# A combination of the geroprotectors trametinib and rapamycin is more effective than either drug alone

**DOI:** 10.1101/2024.07.25.605097

**Authors:** Lisonia Gkioni, Tobias Nespital, Carolina Monzó, Jitin Bali, Taim Nassr, Anna Lena Cremer, Andreas Beyer, Heiko Backes, Sebastian Grönke, Linda Partridge

## Abstract

Genetic suppression of activity of the insulin/IGF/mTORC1/Ras network can ameliorate the effects of ageing in animals. The network provides multiple drug targets because of its role in metabolic disease and cancer, and these are candidates for repurposing for geroprotection. For instance, inhibition of the activity of the mTORC1 complex by rapamycin can extend lifespan in multiple organisms including mice, with early indications of efficacy in humans. Trametinib inhibits MEKs in the Ras pathway and can extend lifespan in *Drosophila*. However, it is not yet known if trametinib alone or in combination with rapamycin can extend mouse lifespan or improve health at older ages. We assessed survival and health indices of female and male mice treated with trametinib or rapamycin alone, or with the two in combination at the same doses. Trametinib treatment extended lifespan in both sexes, while its combination with rapamycin caused further, additive prolongation. Combination treatment reduced liver tumours in both sexes and spleen tumours in males, and ameliorated the age-related increase in brain glucose uptake. There was a striking reduction in inflammation in the brain, kidney, spleen and muscle with combination treatment, accompanied by reduced circulating levels of pro-inflammatory cytokines. Trametinib alone is therefore geroprotective in mice, but combined trametinib and rapamycin treatment is more geroprotective than treatment with either drug alone, suggesting immediate translational potential for humans.

## Introduction

The insulin/IGF/mTORC1/Ras nutrient-sensing network is highly conserved in evolution and is implicated in the aetiology of many age-related diseases ([1]; [2]; [3]). Genetic inhibition of multiple nodes within the network can improve health during ageing and extend lifespan in laboratory model organisms [4], while growing evidence suggests that there is evolutionary conservation of the pro-ageing role of the network in humans [5][2][6]. There is therefore growing interest in the possibility of repurposing existing drugs with targets in this signalling network as geroprotectors to improve human health during ageing [7].

One such example is inhibition of mTORC1 by rapamycin (sirolimus), which is an FDA-approved drug used mainly for immunosuppression after organ transplantation [8] [9]. Rapamycin robustly extends lifespan in multiple model organisms, ranging from worms [10] and flies [11] to mice, where rapamycin administration later in life at 600 days of age increases median and maximal lifespan in both sexes [12]. Rapamycin also ameliorates ageing-related morbidities, including cardiac dysfunction [13] [14] and impaired immune responses [15]. Administration of mTOR inhibitors in a cohort of elderly human subjects led to enhanced immune response to influenza vaccine and reduced infection rate, with briefer and lower dosing than is commonly used clinically and few if any side effects [16, 17]. These findings suggest that some of the geroprotective effects of rapamycin treatment may be, at least partly, conserved in humans.

Reduced signalling through the phosphatidylinositol 3-kinase (PI3K) node of the nutrient-sensing network can extend lifespan in *C. elegans* and *Drosophila* [18] [19], and was for long viewed as the primary route by which the anti-ageing effects of reduced upstream insulin/IGF signalling are mediated. However, Ras signalling plays a role in ageing in yeast [20], while in *Drosophila* the Ras-MEK-ERK pathway is as important a mediator as the PI3K pathway of the effects of reduced upstream insulin/Igf signalling on lifespan [21] [22]. Indirect inhibition of Ras in mice is associated with increased lifespan and enhanced motor function in old age [23]. These findings suggest that inhibition of Ras pathway signalling may have an evolutionarily conserved, geroprotective effect.

Ras hyperactivation is highly oncogenic, with a third of human cancers presenting with a Ras mutation [24], There has therefore been an intense search for small molecule inhibitors that target its activity. Trametinib (also known as Mekinist) is a potent and highly specific small molecule inhibitor of MEK, and is an FDA-approved drug for the treatment of specific melanomas [25]. Oral administration of trametinib increases *Drosophila* lifespan, even when started later in life [22]. However, it is yet to be determined whether the lifespan-extending effects of trametinib are evolutionarily conserved. To examine whether trametinib is geroprotective in mice, we orally dosed female and male mice and assessed their ageing phenotypes.

The insulin/IGF/mTORC1/Ras network is characterised by extensive crosstalk between its branches. It is therefore possible that simultaneous inhibition of different nodes within the network, by combined drug treatments, could be more effective than suppression of single nodes, by prevention of compensatory responses. Combinatorial drug treatments could also induce stronger inhibition of specific targets, or induce a wider range of protective effects, given that different nodes in the network have many non-overlapping targets. Indeed, joint inhibition of mTOR, MEK, and glycogen synthase kinase-3 (GSK-3) by rapamycin, trametinib and lithium, respectively, resulted in additive increases in *Drosophila* lifespan, both with the double treatment combinations and in the triple combination, which yielded a 48% median lifespan extension [26]. However, it is currently unknown whether such combinatorial drug treatment would exert similar benefits in mammalian ageing.

In the present study, we investigated whether administration of trametinib alone or in combination with rapamycin can extend lifespan and improve health at old age in mice. We orally treated male and female mice with trametinib, or rapamycin, or with both drugs at the same doses as in the single drug treatments. We assessed their survival, fitness, brain metabolism and organismal health. Single administration of trametinib or rapamycin significantly increased male and female mouse lifespan, while the combined treatment produced an additive further increase in both sexes. Additionally, the double combination of trametinib and rapamycin significantly reduced liver tumours in both sexes and spleen tumours in males at old age, and alleviated the age-related increased glucose uptake in the brain. Combination treatment also caused a marked reduction of age-related inflammation in brain, kidney, spleen and muscle, accompanied by reduced levels of circulating pro-inflammatory cytokines.

## Results

### Dose range for oral administration of trametinib in mice

To determine the dose range of trametinib that efficiently inhibited Ras-Mek-Erk signalling without adverse effects on health, young, C3B6F1 hybrid, wild type male and female mice were fed with 0.29, 0.58, 1.44, 2.88 or 11.52 mg trametinib / kg of diet for 4 weeks (Fig 1A), and their plasma levels of trametinib (Fig 1B-C), Ras-Mek-Erk pathway activity (Fig 1D-E), body weight (Fig 1F-G) and spleen weight (Fig. 1H-I), Sup. Fig 1) were measured. Trametinib in plasma increased with dietary concentrations and was higher in females than in males (Fig 1B-C). We measured inhibition of MEK activity by western blot analysis on liver samples using phosphorylation of ERK1/2 as read-out (Fig 1D-E). Erk1/2 phosphorylation was unaffected at 0.29 and 0.58 mg/kg trametinib and reduced at concentrations of 1.44 mg/kg trametinib and above in both sexes, although low sample number and high variability meant that only 11.52 mg/kg in female mice resulted in a significant reduction. Trametinib did not significantly affect water uptake (Sup. Fig 1A-B). Trametinib can cause body weight loss, liver lesions and necrosis, and increased alanine aminotransferase (ALT) and alkaline phosphatase (ALP) levels indicative of liver dysfunction [27]. There was no significant effect of any dose of trametinib on plasma levels of AST (Sup. Fig 1C-D) or ALP (Sup. Fig 1E-F). At doses of 0.29-2.88 mg/kg trametinib also did not affect body weight (Fig 1F-G) or spleen size (Fig 1H-I). In contrast, animals fed with 11.52 mg/kg trametinib failed to gain body weight in the 4-week measurement period (Fig 1F-G) with a trend for increased spleen weight in both sexes (Fig 1H-I). In summary, dietary trametinib concentrations of 1.44 mg/kg and above were sufficient to inhibit Ras-Mek-Erk signalling, while only the highest dose of 11.52 mg/kg trametinib induced adverse effects on mouse health. We therefore used 1.44 mg/kg in all subsequent experiments.

**Figure 1.**
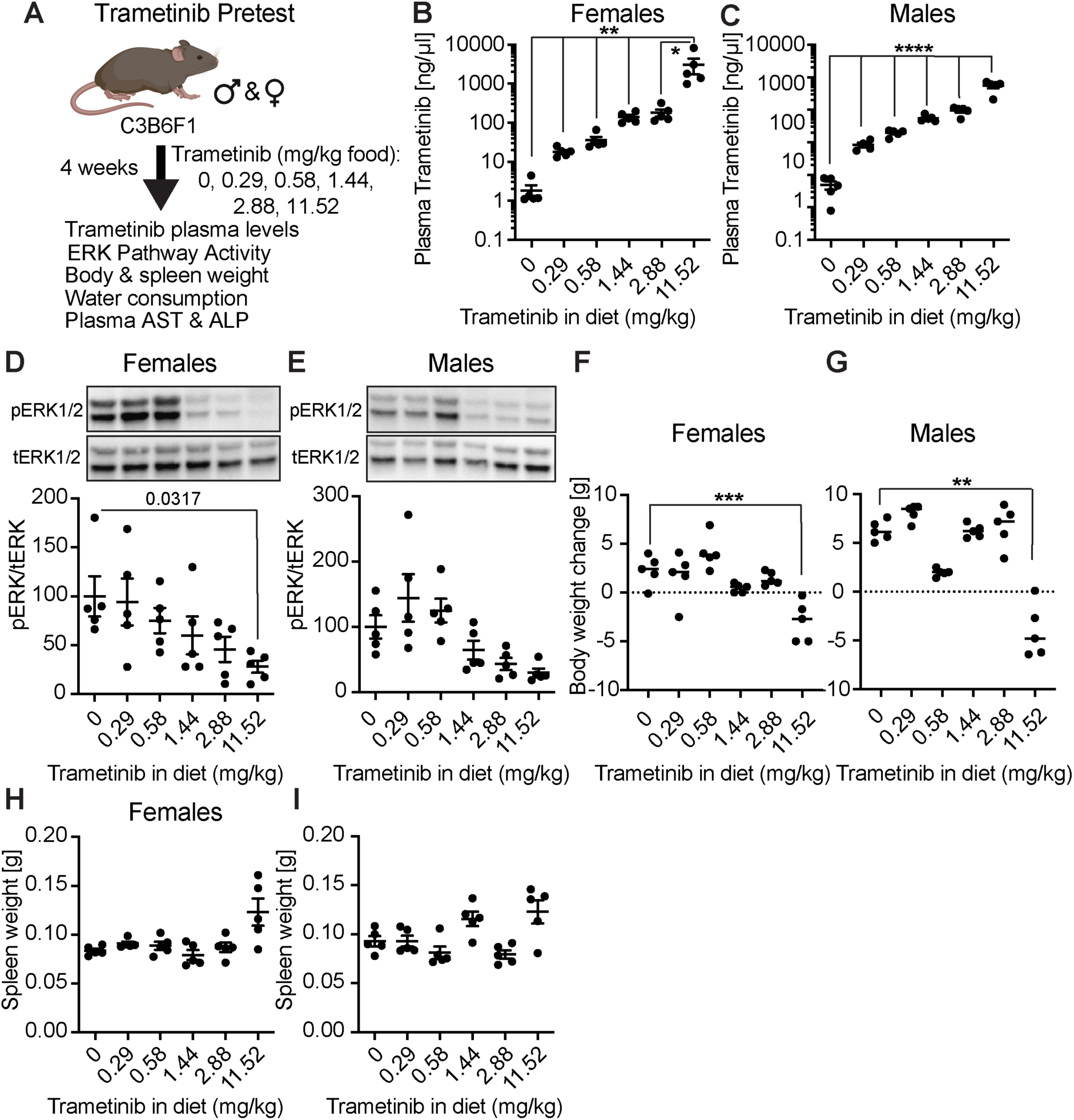
Doses of trametinib that inhibit Ras/MEK/Erk signalling in vivo in mice without detrimental side effects. **a** Schematic outline of the trametinib pretest experiment. Four-month-old C3B6 F1 hybrid mice were used for all experiments. **b-c** Plasma concentration of trametinib in **b** females and **c** males (n=5 per treatment for both sexes, except n=4 for male 0.29) fed for 4 weeks with food containing 0, 0.29, 0.58, 1.44, 2.88 or 11.52 mg/kg of trametinib. Data are presented as mean ± SEM. **d-e** Ras/Mek/Erk pathway activity in liver after 4 weeks of trametinib treatment, by Western blot analysis of Erk1/2 phosphorylation. Representative Western blot (upper panel) and corresponding quantification of **d** female and **e** male liver samples (n=5 per treatment for both sexes). Trametinib concentrations of 1.44 mg/kg and above inhibited Erk1/2 phosphorylation in liver of both sexes. **f**-**g** Change in body weight upon 4 weeks of trametinib treatment in **f** females and **g** males. 11.52 mg/kg trametinib led to a significant reduction in body weight in both sexes compared to control animals (0mg/kg) (n=5 animals per treatment). **h**-**i** Net spleen weight of **h** female and **i** male mice after 4 weeks of trametinib treatment. Spleen weight showed a tendency to increase with 11.52 mg/kg trametinib treatment. One-Way ANOVA with Tukeýs post hoc test: * p<0.05, ** p<0.01, *** p<0.001, **** p<0.001.

### Trametinib extends lifespan and combines additively with rapamycin to further extend

To assess effects of trametinib on survival and health, mice were continuously fed with 1.44 mg/kg of trametinib starting from 6 months of age and their survival measured (Fig. 2A). Trametinib treatment caused a significant lifespan extension in both sexes, in females, with a median lifespan extension of 7.2% (p=0.0384, Log Rank) (Fig 2B) and no significant effect on maximum lifespan (p=0.474, Wang/Allison test), in males with an increase in median lifespan of 10.2% (p<0.0001, Log Rank) (Fig 2C) and maximum lifespan by 15.8% (p=0.014, Wang/Allison test).

**Figure 2.**
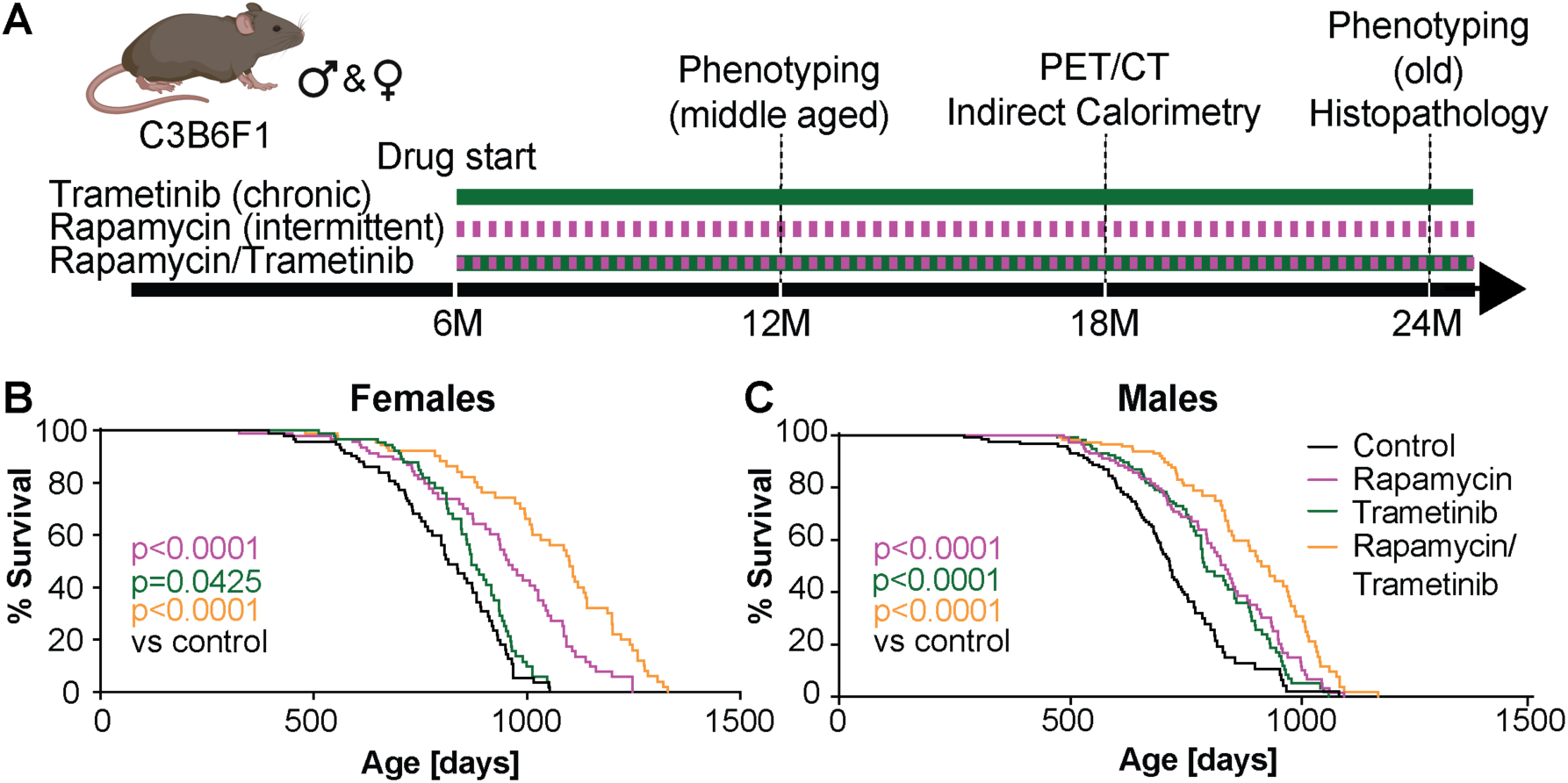
Trametinib alone extends lifespan and acts additively with rapamycin to do so. **a** Schematic overview of trametinib and rapamycin treatment. **b**-**c** Survival of **b** females and **c** males. Trametinib significantly increased survival in both sexes. Combined treatment further increased lifespan. P-values based on log rank test comparing the indicated treatment to control animals. (Sample sizes: Control ♂ n=119, ♀ n=97; rapamycin ♂ n=119, ♀ n=97, trametinib ♂ n=121, ♀ n=97; Combined ♂ n=120, ♀ n=97). Lifespan of control and rapamycin treated animals were previously published [30].

In *Drosophila*, trametinib and rapamycin combine additively to extend fly lifespan [26]. We measured the survival of mice on diets containing either only rapamycin (42 mg/kg) or both rapamycin (42 mg/kg) and trametinib (1.44 mg/kg). 42 mg/kg Rapamycin was chosen as the dose so far inducing the largest increase in longevity [28], and it was fed in alternate weeks (Fig. 2A), which improves metabolic health parameters [15] [29] and extends lifespan to a similar degree as continuous treatment [30]. As previously shown [30], intermittent rapamycin treatment extended lifespan in both sexes (p<0.0001, log rank test) with an increase in median and maximum lifespan of 17.4% and 16.5% respectively in females (Fig. 2B) and 16.6% and 18.3% respectively in males (Fig. 2C). Combined treatment with rapamycin and trametinib increased survival more in females than in males (sex-treatment interaction p=0.0218, Cox Proportional Hazard). Combined treatment caused a larger increase compared to the single treatment (Fig. 2B-C) in both sexes, with median and maximum lifespan increased by 34.9% and 32.4%, respectively, in females (Fig 2B) and by 27.4% and 26.1%, respectively, in males. In females, the hazard ratio of trametinib alone compared to control (p=0.0383) was 0.6926 (0.4886-0.9804), and 0.4478 (0.3002-0.6641) in the presence of rapamycin (p<0.0001).

Rapamycin alone had a hazard ratio of 0.3284 (0.2243-0.4784) compared to controls (p<0.0001), and 0.2123 (0.1380-0.3234) in the presence of trametinib (p<0.0001). In males, trametinib alone had a hazard ratio of 0.5257 (0.3845-0.7169) versus controls (p<0.0001) and 0.5814 (0.4127-0.8154) in the presence of rapamycin (p=0.0017). Rapamycin on its own had a hazard ratio of 0.4623 (0.3376-0.6316) compared to controls (p<0.0001) and 0.5113 (0.3611-0.7208) in the presence of trametinib (p=0.0001). There was thus no indication of a positive or negative interaction between drug treatments in either sex.

In summary, treating mice with both trametinib and rapamycin extended lifespan in both sexes, with lifespan further extended by combined treatment in an additive manner.

### Effects of trametinib and rapamycin on mouse health

We measured general age-related health parameters including body weight (Sup Fig 2A-B), heart function (Sup Fig 2C-F), anxiety behaviour (Sup Fig 2G-J), endurance (Sup Fig 2K-L), motor coordination (Sup Fig 2M-N), and memory (Sup Fig 2O-P) at middle (12 months) and old (20-22 months) age. In addition, we used metabolic cages to measure body weight (Sup Fig 3A-B), fat mass (Sup Fig 3C-D), water and food uptake (Sup Fig 3E-H) and respiratory exchange ratio (RER) (Sup Fig 3I-L) at 16 months of age. Consistent with the pre-test, 1.44mg/kg trametinib had no effect on body weight (Sup Fig 2A-B, Sup Fig 3A-B). In contrast, both rapamycin and combined drug treatment decreased body weight in females at 20 months (Sup Fig 2A, Sup Fig 3A) and in males at 12, 16 and 20 months (Sup Fig 2B, Sup Fig 3B). Combined treatment of females induced a significant increase in fat content, with a similar trend in males (Sup Fig 3C-D). The drug treatments did not affect water or food uptake (Sup Fig 3E-H). Heart rate of both sexes declined with age, and in males both rapamycin and trametinib attenuated the decline, with a similar non-significant trend in females (Sup Fig 2C-D). QT interval, a read-out for left ventricular hypertrophy, increased with age, and rapamycin decreased QT interval in females while both rapamycin and combined treatment prevented the age-related increase in males (Sup Fig 2E-F). The drug treatments had no effect on centre occupancy in the open field arena (Sup Fig 2G-H), forced running endurance on a treadmill (Sup Fig 2K-L) or motor coordination in a rotarod assay (trametinib caused a marginal decrease in females) (Sup Fig 2M-N). Rapamycin-treated females performed slightly worse in the Y maze (Sup Fig 2O), indicating impaired spatial working memory. Noteworthy, rapamycin and combination-treated males showed increased global speed in the Y maze (Sup Fig 2P) with a similar trend for the combined drug treatment in the open field (Sup Fig 3I), suggesting a higher motivation to move at old age. RER was not significantly altered by drug treatment in 16 months old females (Sup Fig 3I,K), while males treated with trametinib, rapamycin or the combination showed significantly reduced RER during the night (Sup Fig 3J), indicating a slight shift in metabolism towards mixed carbohydrate and lipid utilization.

In summary, rapamycin treatment alone or in combination with trametinib reduced body weight in both sexes and attenuated the decline in heart function with age. Both rapamycin and trametinib attenuated the age-related decline in heart rate in males, rapamycin decreased QT interval both sexes and combined treatment prevented the age-related increase in males. Rapamycin- and combination-treated females moved more at late age, and all drug treatments reduced nighttime RER in males.

### Combined drug treatment delays tumour growth

Both trametinib and rapamycin are cancer chemotherapeutics. To address whether their lifespan-extending effects were associated with their anti-cancer function, we performed a cross-sectional histopathological analysis at 24 months of age, complemented by a macropathological assessment of overall tumour load post mortem, to assess any delay in tumour formation. Untreated mice predominantly developed tumours in the liver (Fig 3A-D), spleen (Fig 3E-H) and kidney (Fig 3I-J). Liver tumours were most prevalent, with about 50% of females (Fig 3A) and 70% of males (Fig 3B) with tumours at 24 months of age. No significant differences in liver tumours were detected upon either trametinib or rapamycin treatment at 24 months of age (Fig 3A-B) or post-mortem (Fig 3C-D). However, the combination of rapamycin and trametinib significantly reduced liver tumour numbers in both sexes at 24 months of age (Fig 3A-B) and in males post-mortem (Fig 3D), with a similar trend in females Fig 3C). 25% of females and 20% of males had tumours in the spleen (Fig 3E-F), and 15% of females and 5% of males showed kidney tumours (Fig 3I-J) at 24 months of age. Neither trametinib nor rapamycin reduced tumour formation in the spleen (Fig 3E-F) or the kidney (Fig 3I-J). The combination treatment significantly reduced spleen tumour formation at 24 months of age in males (Fig 3F), with a similar trend in females (Fig 3E). Although we did not detect any tumours in the kidney upon combined treatment, this tumour type was so rare in controls that the effect was not significant.

**Figure 3.**
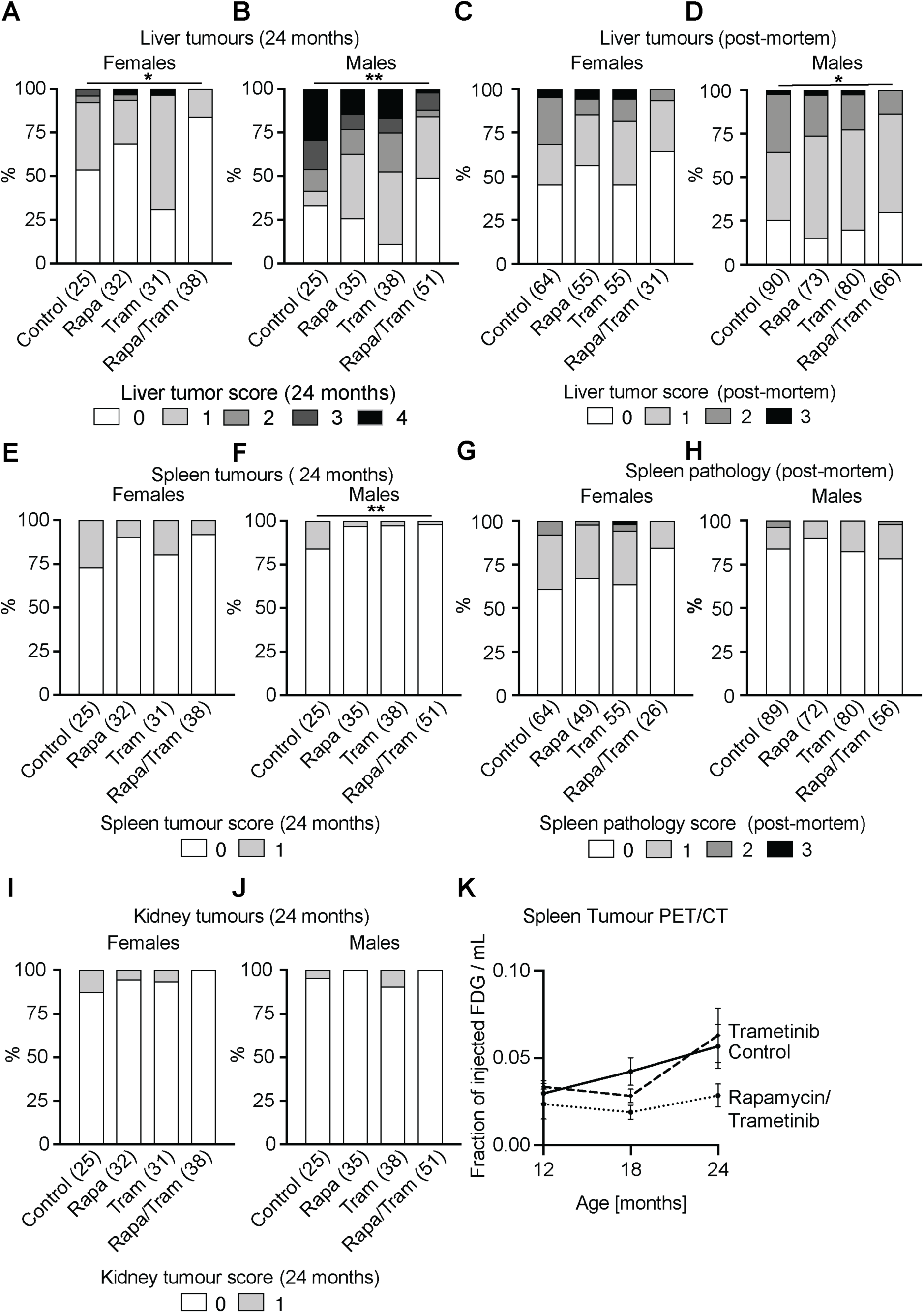
Trametinib and rapamycin combined treatment delays tumour formation and overall tumour load. Cross sectional histopathological analysis of tumour levels in **a**-**b** liver, **e**-**f** spleen and **i**-**j** kidney samples of 24 months old females (**a**, **e**, **i**) and males (**b**, **f**, **j**). Combined drug treatment significantly reduced liver tumours in both sexes at 24 months and spleen tumours in males, with a similar trend for reduced tumour load in female spleen and male and female kidney (**e**-**f**). The absence of liver tumours was scored with 0 and the presence of low, moderate, high and very high severity tumours was scored with 1, 2, 3 and 4, respectively. The absence or presence of spleen and kidney tumours was scored with 0 or 1, respectively. Liver tumour data for rapamycin and control animals were previously published [30]. Post-mortem macropathology analysis of **c**-**d** liver and **g**-**h** spleen of animals from the lifespan and phenotyping cohort. **k** Longitudinal analysis of spleen tumour progression via 18-FDG PET/CT measurements in the same female animals at 12, 18 and 24 months of age. While tumour load increased from 12-24 months in control and trametinib-treated mice, combination-treated animals showed no increase. Data are presented as mean ± SEM. 18-FDG uptake of Rapamycin/Trametinib versus control at 24 months p-value= 0.08. Two-Way ANOVA with post hoc Bonferroni test and simple linear regression. (Control n=7, Trametinib n=5 and Rapamycin/Trametinib n=4). Data in **a**-**j** are presented as percentage over total and statistical analysis was performed by Chi-square test and Poisson regression. *p<0.05, **p<0.01. Numbers in brackets indicate the number of scored tissues per treatment.

In summary, only the combination of rapamycin and trametinib caused significant effects on tumour growth and this was more obvious in the cross-sectional analysis, e.g. compare spleen pathology in males at 24 months and post-mortem (Fig 3F vs 3G), which may indicate a delay in tumour formation rather than reduced overall lifetime tumour load. To investigate this further, we performed a longitudinal analysis, measuring tumour formation *in vivo* by fludeoxyglucose F18 (18F-FDG) positron emission tomography (PET) coupled with computed tomography (CT) [31] in the same animals at 12, 18 and 24 months of age (Fig 3K). The capacity for the PET/CT measurement was limited, and we therefore restricted our analysis to control, trametinib and combination-treated females. Tumours were identified as hypermetabolic regions, first in 24 months old animals and then traced back in the same animals at 18 and 12 months of age. 18F-FDG PET/CT identified tumours in several organs, including the liver and uterus, but we were only able to reliably quantify tumour progression in the spleen (Fig 3G). Tumour occurrence increased with age in control animals and, consistent with the histopathological analysis (Fig 3E), trametinib had no effect on spleen tumours at 12, 18 or 24 months (Fig 3K). In contrast, combination-treated animals showed lower spleen tumours at all 3 ages, with no significant increase with age. These results are consistent with the hypothesis that the combined drug treatment delayed tumour formation but, due to the missing single rapamycin treatment, it is possible that the effect was attributable to rapamycin.

### Combined treatment with rapamycin and trametinib attenuates age-related increase in brain glucose uptake

Glucose uptake in the brain increases during ageing in mice [32] and may be linked to impaired cognitive function [33]). We therefore used the 18F-FDG PET/CT longitudinal data to address whether trametinib and combined drug treatment could attenuate the age-related increase in glucose uptake (Fig 4A-G). As expected, 18F-FDG uptake increased with age in control females across the whole brain (Fig 4A, E), consistent with a previous study [32]. There was a significant increase in 18F-FDG uptake between 12 and 24 months in the striatum and cerebellum, but not the cortex (Fig 4A, E). Females treated with trametinib showed a significant increase in 18F-FDG uptake across the whole brain and in the striatum, cerebellum and cortex between 12 and 18 months of age (Fig 4B, F), with no further increase between 18 and 24 months. Combined treatment with rapamycin and trametinib protected animals from the age-related increase in glucose uptake both across the whole brain and in the striatum, cortex and cerebellum (Fig 4C, G). Two-way ANOVA analysis confirmed that age-related glucose uptake was significantly different between control and drug-treated animals (p<0.05 interaction between age and drug treatment, Fig 4D). Direct comparisons between changes in 18F-FDG uptake between 12 and 24-months (Fig 4F-H) showed that trametinib-treated animals had an intermediate phenotype between control and combination-treated animals with a clear age-related increase in glucose uptake, while there was no change in the double drug-treated mice. These results suggest that trametinib treatment ameliorated the age-related increase in brain glucose uptake, but that it only completely blocked it when combined with rapamycin.

**Figure 4.**
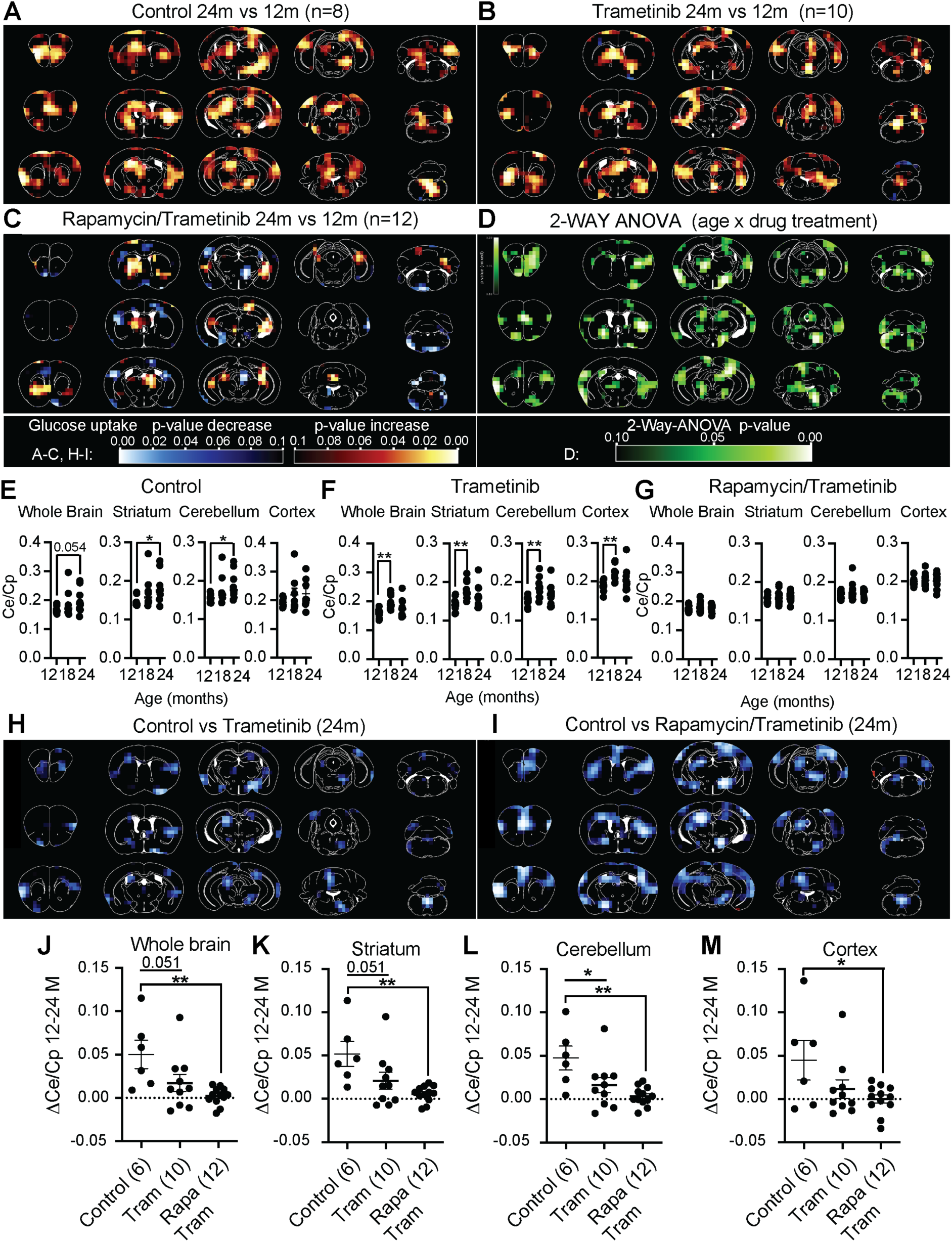
Combined treatment with trametinib and rapamycin counteracts the age-related increase in brain glucose uptake. Longitudinal 18-FDG PET/CT measurements of age-related changes in differential glucose uptake in the female mouse brain. **a**-**c** Images show differential regional glucose uptake across the mouse brain between 24 and 12 months old **a** control, **b** Trametinib and **c** Rapamcin/Trametinib treated female mice. Colour code represents the p value for the indicated voxels in a paired Student’s t test comparing 24 to 12 months old animals. Increase in glucose uptake is shown in red/yellow, decrease in blue colour. Numbers in brackets indicate the number of animals analysed per treatment. **d** Two-Way ANOVA analysis highlights brain regions that showed a significant interaction between age and drug treatment in differential glucose uptake. A lighter green colour indicates a higher significance in the 2-Way ANOVA analysis. **e**-**f**: Quantification of glucose uptake in whole brain, striatum, cerebellum and cortex at 12, 18 and 24 months of age in **e** control, **f** trametinib and **g** rapamycin/trametinib-treated mice. Glucose uptake significantly increased with age in striatum and cerebellum of control animals, but not in combination-treated animals. Data are presented as mean ± SEM. *p<0.05, **p<0.01. **h**-**i** Images show differential glucose uptake in the brain of 24 months old **h** trametinib vs control and **i** combination vs control mice. **j**-**m** Quantification of differential glucose uptake between 12 months and 24 months old **j** whole brain, **k** striatum, **l** cerebellum and **m** cortex. Decrease in glucose uptake is indicated in blue colour. Age-related differences in glucose uptake between 12 and 24 months were significantly lower in combination-treated animals compared to controls. A similar but milder effect was also observed in trametinib-treated animals, which was only significant in the cerebellum. *p<0.05, **p<0.01. Two-Way ANOVA with Bonferroni post hoc test. Data are presented as median with SEM.

### Combined treatment with trametinib and rapamycin reduces activated microglia and astrocytes in old mouse brain

Brain inflammation increases the requirements for glucose and results in greater 18F-FDG uptake [34] [35]. Microglia are the primary immune cells of the brain and their increased activity has been associated with age-related brain inflammation accompanied by synapse loss and late-onset neurodegeneration [36] [37]. Microglia act in concert with astrocytes to resolve inflammation and astrocyte-dependent Tgfβ secretion can inhibit microglia activation [38]. Ageing disrupts this interplay leading to exacerbated inflammation in both an autonomous and astrocyte-dependent manner [39][40]. Since Ras-MEK-Erk signalling has been implicated in microglial activation [36], we assessed whether drug treatment reduced activation of microglia and astrocytes by performing immunohistological stainings of brain sections using antibodies against Iba-1 and Gfap for activated microglia and astrocytes, respectively (Fig 5A-E) in brains of 24-month-old female control (Fig 5B), rapamycin (Fig 5C), trametinib (Fig 5D) and rapamycin/trametinib (Fig 5E) treated mice. To address age-related changes, we also stained brains of 6-month-old control mice (Fig 5A). Consistent with the age-related increase in glucose uptake, activated microglial and astrocyte density significantly increased in the striatum (Fig 5F-G), hippocampus (Sup Fig 4A-B) and cortex (Sup Fig 4C-D), but not the cerebellum (Sup Fig 4E-F) of 24-month compared to 6-month-old control mice, indicating increased brain inflammation at older age. Combined drug treatment significantly reduced the density of activated microglia and astrocytes in the striatum of 24-month-old mice (Fig 5F-G), with a similar trend in the cortex (Sup Fig 4C-D), but no significant effects of the double drug treatment in the hippocampus (Sup Fig 4A-B) or the cerebellum (Sup Fig 4E-FE-F). Compared to the combined treatment, single treatment with rapamycin or trametinib was not as efficient in reducing activated microglia density in the striatum (Fig 5F). In summary, both rapamycin and trametinib had mild beneficial effects, but only the combination of both drugs reduced age-related brain inflammation in a region-specific manner. Noteworthy, unlike microglia density, 18F-FDG uptake was also increased in the cerebellum, suggesting that mechanisms other than brain inflammation may have contributed to the increase.

**Figure 5.**
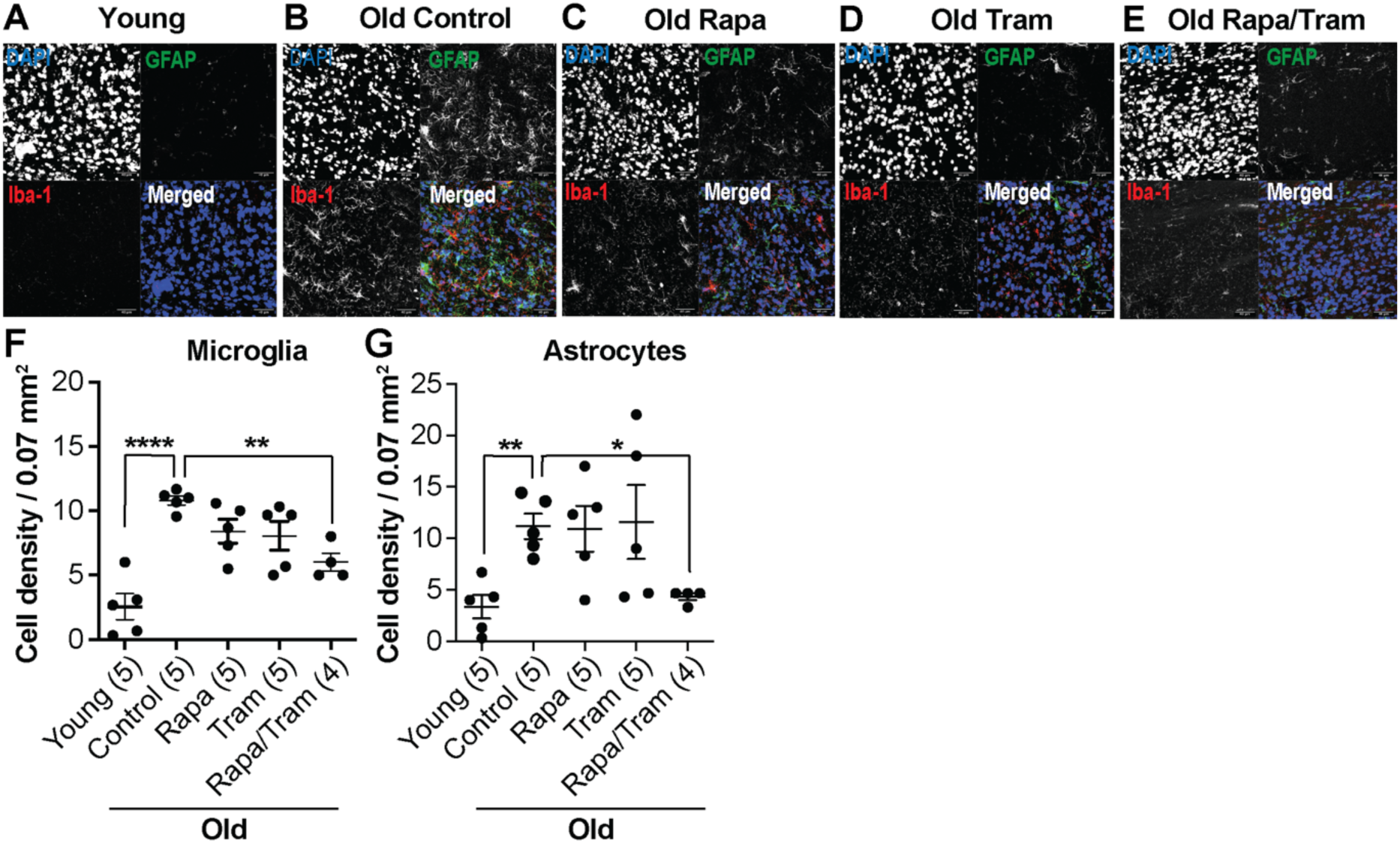
Combination treatment attenuates age-related brain inflammation in the striatum. **a-e** Representative confocal composite images of immunohistochemical stainings of the striatum region in **a** 6-month-old young controls, **b-e** 24-month-old: **b** controls, **c** Rapamycin-treated, **d** Trametinib-treated and **e** Rapamycin/Trametinib-treated female mice. Nuclei were stained with DAPI, astrocytes with GFAP and activated microglia with Iba-1. **f**-**g** Quantification of **f** microglia density (DAPI/Iba-1 positive cells) and **g** astrocyte density (DAPI/GFAP positive cells) per mouse striatum of young and old control animals and old drug-treated animals. Microglia and astrocyte density significantly increased with age in the striatum of control animals. Combination treatment significantly reduced microglia and astrocyte density in the striatum compared to old controls. *p<0.05, **p<0.01, ****p<0.0001. One-Way ANOVA with Bonferroni post hoc test. 10-15 confocal images per brain region were averaged per sample. Numbers in brackets indicate the n number per treatment. Data are presented as mean ± SEM.

As Ras-MEK-Erk signalling has directly been implicated in microglia activation [36], we next addressed whether trametinib was able to cross the blood-brain barrier. Mass spectrometry was used to measure trametinib levels in the brains of the animals used for the trametinib pre-test (Sup Fig 4G-H). At dietary concentrations of 0.29 and 0.58 mg/kg, trametinib was not detected in the brain (Sup Fig 4G-H). Significant, but very low, levels of trametinib were detected, with levels in the brain 25- and 30-fold lower than the corresponding plasma concentrations of female (140 ng/µl) and male (55,8 ng/µl) mice, respectively. This result suggests that trametinib supplied orally with 1.44mg/kg barely crosses the blood brain barrier, consistent with previous observations that the brain distribution of trametinib is restricted by active efflux at the blood brain barrier [41]. Only at 11.52 mg/kg, the highest concentration used in the pre-test, did we detect trametinib in the brain, which might indicate saturation of the efflux system due to the high trametinib plasma levels. In conclusion, the effects on microglia upon treatment with 1.44 mg/kg trametinib might be indirect and caused by the action of trametinib in peripheral tissues.

### Combined treatment with trametinib and rapamycin reduces inflammation in peripheral tissues

Ageing is characterised by increased levels of inflammation in multiple peripheral organs, which manifest in a low-level systemic pro-inflammatory phenotype, termed ‘inflammaging’ [42]. Inflammaging is associated with both mTOR and Erk pathway activity [43] [44] [45] [45, 46]. As the combined treatment with rapamycin and trametinib reduced brain inflammation, we next assessed inflammation in peripheral tissues, by histopathology of kidney and white adipose tissue (WAT) (Fig 6A-D), which are affected by age-related inflammation [47] [48] [49]. Control animals showed high levels of kidney inflammation with 64% of females and 60% of males affected at 24 months of age (Fig 6A-B). While single treatment with trametinib did not reduce kidney inflammation (Fig 6A-B), rapamycin induced a non-significant reduction, with only 40% of females and 43% of males affected at 24 months. Combined treatment with rapamycin and trametinib significantly reduced kidney inflammation in both sexes with only 34% of females and 35% of males affected. These findings suggest that most of the combined drug effect was due to rapamycin, which positively affects kidney health [50] [51]. Control mice also showed high levels of inflammation in the WAT, with 82% of females and 86% of males affected (Fig 6 C-D). No drug treatment significantly reduced WAT inflammation, with 69%, 63% and 65% of females and 81%, 76% and 65% of males affected in trametinib-, rapamycin- and combination-treated animals, respectively.

**Figure 6.**
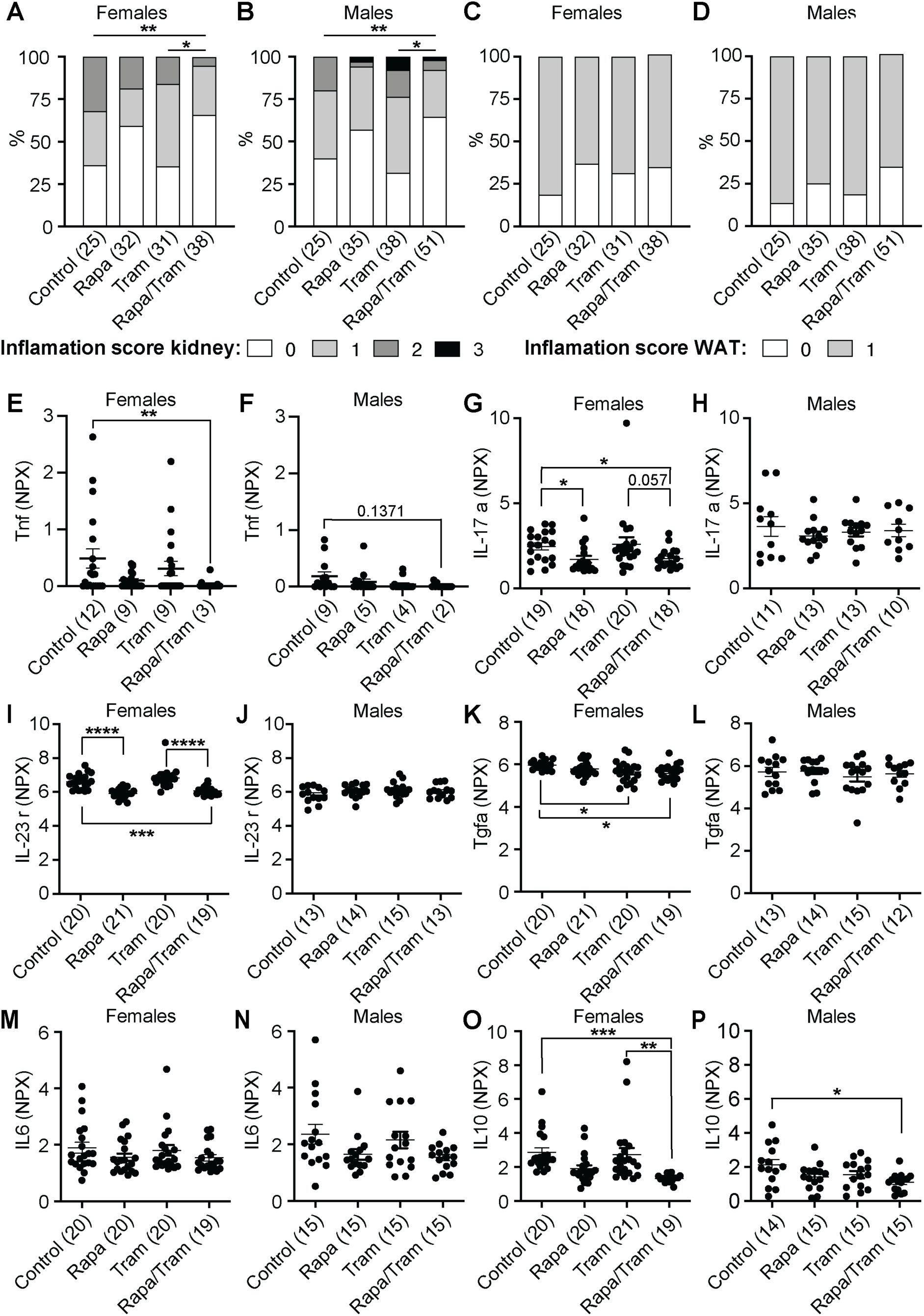
Combination drug treatment reduces peripheral inflammation. **a-d** Histopathological analysis of inflammation in **a-b** kidney and **c-d** white adipose tissue (WAT) of 24 months old females (**a**, **c**) and males (**b**, **d**). Combination-treated females and males showed significantly reduced kidney inflammation compared to control and Trametinib-treated animals. Linear regression. *p<0.05, **p<0.01. There was a non-significant trend for reduced WAT inflammation upon treatment with Rapamycin and Rapamycin/Trametinib. **e**-**p** Plasma protein levels of circulating cytokines as measured by the Olink Target 96 Mouse Exploratory panel. **e-f** Tnf, **g-h** IL17a, **i-j** IL23r, **k-l** Tgfα, **m-n** IL6 and **o-p** IL10. Drug effects on plasma cytokine levels were highly sex-specific, with Tnf, IL-17a, IL23r, Tgfa being downregulated by the combination treatment only in females but not males. *p<0.05, ***p<0.001, ****p<0.0001 One-Way ANOVA with Tukey post hoc test. Numbers in brackets indicate the number of plasma samples from independent mice per treatment. Inflammation and plasma cytokine data for control and rapamycin treated animals were previously published [30].

We also assessed inflammatory markers in plasma of both sexes at 24-months, using the Olink Target 96 Mouse Exploratory panel (Fig 6E-L, Sup Table 1). Consistent with the effect on tissue inflammation, combination-treated females showed significantly reduced plasma levels of key pro-inflammatory proteins [52], including Tumour necrosis factor (Tnf, Fig 6E), Interleukin-17a (IL17a, Fig 6G), Interleukin-23r (IL 23r, Fig 6I), and Transforming growth factor *α* (Tgfa, Fig 6K). Trametinib-treated females showed a significant reduction only in Tgfa (Fig 6K), while rapamycin treatment induced significantly reduced levels of Interleukin-17a (IL-17a, Fig 6G) and Interleukin-23r (IL-23r, Fig 6I). The single drug treatments had no significant effect on pro-inflammatory cytokines in males, with only IL6, which was not regulated in females (Fig 6M), showing a trend to be downregulated in males (Fig 6N). Combination-treated males showed a trend for reduced Tnf (Fig 6F) and IL6 levels (Fig 6N). Thus, combination treatment induced a sex-specific reduction in females of circulating pro-inflammatory cytokines. Noteworthy, combination treatment significantly reduced the level of the anti-inflammatory cytokine Interleukin 10 (IL-10) in both females (Fig 6O) and males (Fig 6P), consistent with previous findings [53]. In summary, our findings indicate that combination treatment reduced inflammation in peripheral tissues accompanied by reduced levels of circulating pro-inflammatory cytokines.

### Combined treatment with rapamycin and trametinib ameliorates spleen pathology but not rapamycin-induced liver lipidosis and testicular lesions

We assessed the impact of the drug treatments on age-related, non-neoplastic pathologies, by measuring heart hypertrophy, kidney and spleen pathology at 24 months of age (Sup Fig 5). Heart hypertrophy and kidney glomerulopatholgy were more pronounced in male than female mice, but there was no significant effect of any of the drug treatments (Sup Fig 5A-D). Trametinib had no significant effect on spleen pathology in females and caused a mild non-significant reduction in males (Sup Fig 5E-F). Females treated with rapamycin showed a trend for reduced spleen pathology with a significant reduction in males (Sup Fig 5E-F). Combined treatment of rapamycin and trametinib significantly reduced spleen pathology in both sexes (Sup Fig 5E-F). Increased spleen size in old mice reflects disrupted microarchitecture, extramedullary hematopoiesis, chronic inflammation, and accumulation of senescent cells [54] [55], all of which impair spleen function. Interestingly, spleen weight was significantly decreased by trametinib in females, with a similar trend in males and by rapamycin and combination treatment in both sexes (Sup Fig 5G-H), suggesting that both drugs combat spleen immunosenescence.

Long-term administration of rapamycin is associated with negative side effects, including liver lipidosis and testicular degeneration [56] [57]. In contrast to rapamycin, trametinib treatment did not induce liver lipidosis or gonadal lesions (Sup Fig 5I-L), while animals treated with both drugs showed similar increases in liver lipidosis and testicular degeneration as rapamycin-treated mice (Sup Fig 5I-L). Thus, trametinib treatment did not induce detrimental side effects as from rapamycin, but nor did it protect rapamycin-treated mice from these pathologies.

### The transcriptome responds to drug treatment in a tissue-, drug- and sex-specific manner

Combination drug treatment resulted in reduced age-related kidney inflammation and spleen size. Neutrophil infiltration into ageing muscle also occurs in ageing mice [58]. We therefore measured gene expression changes in response to drug treatment in kidney, spleen and muscle of 24 months mice using RNA Seq to gain further insight into mechanisms (Fig 7, Sup Table 2). In all three organs, the most strongly regulated genes were linked to inflammatory processes (Figure 8, Sup Fig 6-7).

**Figure 7.**
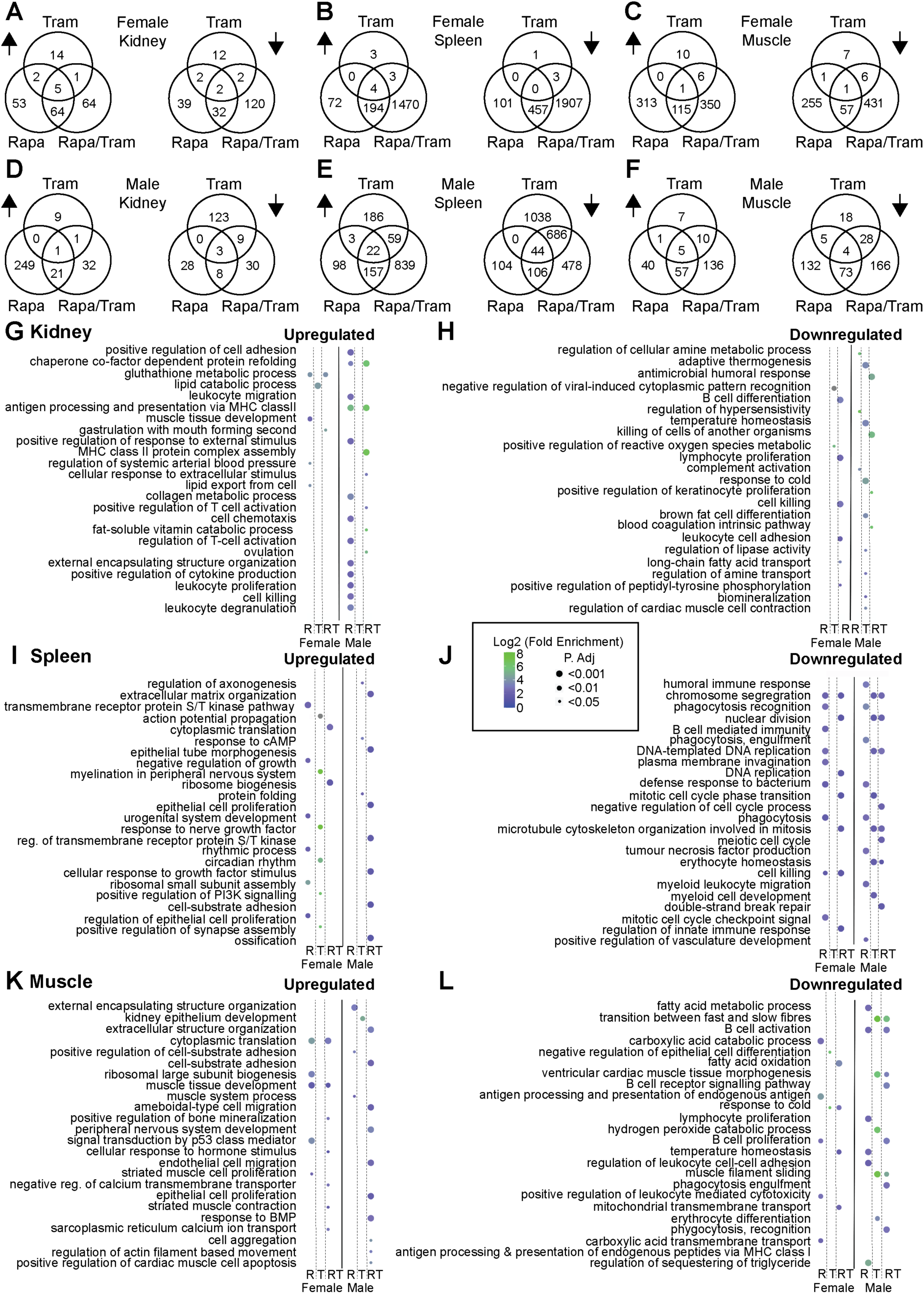
The transcriptome responds in a highly tissue-, drug- and sex-specific manner towards rapamycin and trametinib treatment. RNA seq analysis of kidney, spleen and muscle tissues isolated from 24 months old male and female mice used to measure gene expression changes in response to Rapamycin, Trametinib and Rapamycin/Trametinib treatment. **a-f** Venn diagrams depicting the overlap in differential gene expression upon Rapamycin, Trametinib and Rapamycin/Trametinib treatment in **a**, **d** kidney, **b**, **e** spleen and **c**, **f** muscle tissue. The left Venn diagram per panel shows the overlap in upregulated genes (upward arrow) and the right panel shows downregulated genes (downward arrow). **g**-**l** Gene ontology (GO) term analysis of significantly differential expressed genes upon Rapamycin, Trametinib and Rapamycin/Trametinib treatment. Dotplot representation of significantly enriched top GO terms sorted by adjusted *p*-value for upregulated (**g**, **l**, **k**) and downregulated (**h**, **j**, **l**) genes in **g**, **h** kidney, **i**, **j** spleen and **k**, **l** muscle. The colour of the dots indicates log2 fold enrichment, the size of the dots the adjusted p value for enrichment of the specific GO term. *n* = 4-5 per drug treatment.

The number of differentially expressed genes (DEGs) was strongly dependent on drug treatment and tissue type (Fig 7A-F). Trametinib treatment caused only minor changes in the transcriptome of female and male kidney (40/146 DEG) and muscle (32/78 DEG) and the female spleen (14 DEG), while in male spleen 2038 genes were differentially regulated. Apart from in the male spleen, more genes were differentially regulated upon rapamycin treatment, with 199/210 DEGs in kidney, 828/534 DEGs in spleen and 743/317 DEG in muscle of female and males, respectively. The strongest changes in gene expression occurred with combination treatment, with 290/105 DEGs in kidney, 4038/2391 DEG in spleen and 967/479 DEGs in muscle of female and males, respectively. Gene ontology (GO) enrichment analysis showed little overlap in the transcriptional responses in males and females (Fig 7G-L). Only GO terms associated with chromosome segregation, nuclear division and microtubule cytoskeleton organization involved in mitosis were downregulated in the spleen of both male and female - combination treated animals (Fig 7J). There was also little overlap in DEGs between rapamycin and trametinib (Fig 7A-F), with gene ontology (GO) analysis showing different terms between rapamycin and trametinib single treatments (Fig 7G-L). Although there was significant overlap in DEGs between the single and the combined drug treatments (Fig 7A-F), most DEG were specific to the combined treatment, and the GO term analysis also showed little overlap in GO terms between the single and combined treatments (Fig 7G-L), suggesting that the stronger effect in the combined treatment was not simply a result of the combination of the individual drug effects. Many GO terms were associated with immune regulation and inflammation (Fig 7G-L). In the kidney these were found both among the up and downregulated genes (Fig 7G-H). In contrast, in the spleen and muscle genes associated with immune regulation and inflammation were exclusively downregulated (Fig 7I-L), suggesting reduced inflammaging in these tissues.

Based on the GO term analysis we also looked for genes associated with inflammation. Cd5l and Chemokine (C-C motif) ligand 8 (Ccl8, also known as Mcp2) were among the strongest downregulated genes by the combined treatment in the kidney of female and male mice and were also downregulated in male and female spleen and male muscle (Sup Fig 6A-F). Increased levels of Cd5l and Ccl8 have been linked to inflammatory conditions [59] and invasiveness and metastasis in various cancers [60]. Cd5l and Ccl8 were also downregulated by rapamycin but not trametinib treatment in the spleen (Sup Fig 6B, 6E) and in tendency also in the kidney (Sup Fig 6A, 6D). Reduced expression of CD5L, which is expressed mostly by macrophages of inflamed tissues [59], suggests reduced macrophage infiltration of kidney tissue of combination and rapamycin treated animals. Lysozyme 1 (Lyz1) expression was strongly downregulated by the combination treatment in both kidney and spleen of male and female mice (Sup Fig 6G-H) but was not regulated in muscle. Lyz1 has a pro-inflammatory function and can promote kidney fibrosis upon overexpression [61]. In the kidney Lyz1 was downregulated by trametinib but not rapamycin treatment (Sup Fig 6G). In the female spleen Lyz1 was significantly downregulated by rapamycin and trametinib, while single drug treatments did not reduce Lyz1 levels in the male spleen. Reduced Lyz1 levels upon trametinib treatment might suggest a protective role of trametinib against kidney fibrosis, which is even more pronounced after coadministration of rapamycin. Combined drug treatment also reduced expression of chemokines and chemoattractants in the kidney, spleen, and muscle, including the Chemokine (C-X-C motif) ligand (CXCL) proteins Cxcl7, Cxcl11 and Cxcl13 (Sup Fig 6I-L). Cxcl7 was downregulated by rapamycin in male muscle and by the combined treatment in male muscle and kidney (Sup Fig 6I-J). Cxcl11 was significantly downregulated by rapamycin and the combination treatment in female spleen (Sup Fig 6K). Cxcl13 was in the top 10 of significantly downregulated genes by rapamycin in muscle, with a similar trend for the combination treatment (Sup Fig 6L). Noteworthy, Cxcl14 was significantly upregulated by rapamycin and the combination treatment in muscles of both sexes (Sup Fig 6M). In contrast, Cxcl14 was significantly downregulated in the female but not male spleen under combination treatment, (Sup Fig 6N). CXCL14 has context-dependent pro- and anti-tumor effects, which could explain the tissue specific regulation [62]. Cxcl7, Cxcl11 and Cxcl13 act as activators of neutrophils and macrophages, thereby promoting inflammatory states [63], while also facilitating tumour growth, invasion and migration [64][65]. Thus, their reduced expression upon combined drug treatment is consistent with the anti-inflammatory and anti-cancer effect of rapamycin and trametinib co-administration.

Ets transcription factors are key downstream effectors of the Ras/MEK/Erk pathway and activated Ets2 plays a crucial role in sustaining inflammatory responses by regulating the expression of inflammatory genes, such chemokines and matrix metalloproteinases (Mmp) [66] [67]. Spleen tyrosine kinase (Syk), a direct target of Ets2 (Ghosh et al., 2012), was significantly downregulated by combination but not single drug treatment in female kidney (Sup Fig 7A). Syk was also downregulated by rapamycin and the combination treatment in male muscle tissue (Sup Fig 7B). SYK deficiency has been shown to be protective against inflammation [67], suggesting that downregulation of Syk contributes to reduced inflammation in combination treated mice. Expression of the two matrix metalloproteinases Mmp2 and Mmp13 was significantly downregulated in the female but not the male spleen upon combination treatment, and with rapamycin for Mmp2 in the female spleen (Sup Fig 7C-D). MMPs are involved in inflammation and increased Mmp13 and Mmp2 expression has been associated with increased tumour invasion and cancer progression [68][69]. Mmp2 interacts with the α-2-macroglobulin (A2m) gene, which was among the strongest downregulated genes by rapamycin and combination treatment in the spleen (Sup Fig 7E). A2m modulates immune and inflammatory processes [70], and promotes pro-proliferative and anti-apoptotic responses in cancer cells and macrophages [71]. A2m expression was also strongly downregulated by rapamycin and the combination treatment in the female kidney ((Sup Fig 7F). Elevated A2m levels have been associated with several kidney pathologies including fibrosis and age-related kidney dysfunction [72]. Thus, rapamycin and the combination treatment might alleviate age-related kidney fibrosis and dysfunction by lowering A2m levels. Interestingly, levels of Mmp12 were strongly increased by rapamycin and the combination treatment in both the female and male spleen (Sup Fig 7G). Mmp12 usually promotes inflammation, but has also been associated with anti-inflammatory effects [73]. Inflammatory gene expression was often sex-specific. While the drug treatments only reduced Mmp expression ín females, levels of chymotrypsin-like elastase (Cela) genes were exclusively downregulated in male mice (Sup Fig H-J). Cela2a and Cela3b levels were significantly downregulated by single and combination drug treatments in the male but not female spleen (Sup Fig7 H-I). Furthermore, Cela1 was downregulated by rapamycin in the male muscle (Sup Fig7 J). Cela genes are associated with inflammatory states [74] and are also upregulated in cancer [75]. In summary, the treatment with rapamycin and trametinib induced a drug, sex and tissue specific remodelling of the ageing transcriptome characterised by reduced expression of genes associated with inflammation and cancer growth.

## Discussion

In this study, we establish the FDA-approved drug trametinib, an inhibitor of RAS-Mek-Erk signalling, as a gero-protective drug in mammals. We further show that the combination of trametinib and the mTOR inhibitor rapamycin produces an even greater lifespan extension compared to the single drug treatments in both female and male mice. The combination treatment attenuated the decline in heart function with age, delayed tumour growth and overall tumour load, and reduced brain and peripheral inflammation, suggesting improved health at old age. Several ongoing clinical trials address the potential of mTOR inhibitors as geroprotective drugs in humans [76]. Our study suggests that simultaneous inhibition of the mTOR and Ras-Mek-ERK pathway provides additional benefits that are worth exploring in humans.

In patients, trametinib is used for the treatment of metastatic melanomas [76] and has also been used in clinical trials for the treatment of solid tumours [77] [78]. At a dose of 2 mg trametinib once a day, the drug was well tolerated with only small side effects including skin rash, diarrhoea, fatigue and retinopathy [77]. Average peak plasma levels in patients receiving 2mg trametinib daily were between 5.5 and 7.5ng/ml [77] [78], which is significantly higher than the 0.1ng/ml in mice fed with the lifespan extending dose of 1.44mg trametinib/kg food. In a mouse model of mucosal melanoma, administering trametinib via oral gavage resulted in an average peak concentration of 23.8ng/ml [79], which is more than 200-fold the concentration required for lifespan extension. Although trametinib plasma levels might not be directly comparable between chronic feeding and acute application, these results suggest that higher trametinib doses are required for its anti-cancer effect than for lifespan extension. Consistently, tumour progression or load were not significantly changed in long-lived mice treated with trametinib. The effect of trametinib at the chosen concentration of 1.44mg/kg on lifespan and health at old age was mild, indicating that we underdosed and that the optimal dose for lifespan extension might be higher. In the pre-test, a dose of 2.88 mg/kg trametinib did not cause detrimental side effects. Thus, this can be used as a guideline for future studies to optimize the dose of trametinib for maximal lifespan extension. Additionally, it would be interesting to determine whether late-onset or intermittent feeding recapitulates the effects of continuous trametinib treatment on mouse lifespan, as this would be key to enhance adherence and feasibility of trametinib treatment in humans.

Combined treatment with rapamycin and trametinib increased lifespan by 29% and 27% in female and male mice, respectively. There was no indication of any positive or negative interaction between the two drugs, which had an additive effect on lifespan. There was a similar additive effect of these two drugs on lifespan in *Drosophila*, but there the doses of each that were used were the ones that alone had produced a maximum increase. It is not known what doses of rapamycin or trametinib produce a maximum increase in lifespan in mice, or if these would also maximise late-life health. The combination of rapamycin with metformin increased mouse lifespan by 26% and 23% in females and males, respectively [80], while coadministration of rapamycin and acarbose led to a 28% and 34% increase in median lifespan in females and males, respectively [29, 80]. Thus, the different drug combinations have similar effects on mouse lifespan and it might be interesting to combine more than two drugs simultaneously to test whether they act additively. In *Drosophila*, the triple combination of rapamycin, trametinib and lithium further increased lifespan compared to single or double treatments [26]. In contrast to flies, lithium treatment does not extend lifespan in mice [81] and was therefore not included in the current study.

In contrast to rapamycin, which caused liver lipidosis and testicular degeneration, trametinib treated mice showed no obvious detrimental side effects, demonstrating its safe use, at least in mice. Apart from their longer lifespan, trametinib treated mice only showed mild phenotypes and the mechanisms contributing to their increased lifespan are currently unclear. Interestingly, trametinib treatment reduced spleen size at old age and caused a strong transcriptional response especially in the male spleen with an enrichment of GO terms associated with cell division and myeloid cell development. The Ras-Mek-Erk pathway controls B and T cell development [82] [83] and Erk controls hyperproliferation in a mouse model of lymphoproliferative disease characterised by an enlargement of peripheral lymphoid organs [83]. Thus, trametinib might affect longevity by affecting immunosenescence in the ageing spleen, a hypothesis that should be addressed in future experiments.

The combination treatment with rapamycin and trametinib ameliorated the age-related increase in brain glucose uptake and reduced density of activated microglia and astrocytes in several brain regions of female mice, most prominently in the striatum, indicative of reduced brain inflammation in double treated animals. Since the drug did not cross the blood-brain barrier at the dose used, this finding implies that something else does so to decrease age-related inflammation in the brain. Treatment with the pro-longevity drugs acarbose, 17-α-estradiol and nordihydroguaiaretic acid specifically reduced age-associated inflammation in the hypothalamus of male but not female mice, while caloric restriction had positive effects on brain inflammation in both sexes [84]. Thus, since effects of geroprotective drugs can be sex specific and microglia show sex related differences during development and ageing [85], it will be important for future studies to test whether the combination treatment is also protective against brain inflammation in male mice. Another limitation of our study is, that we did not measure brain glucose uptake in rapamycin treated mice, and we can therefore not address whether the changes observed in combination treated animals are due to a combinatorial effect or just caused by rapamycin. The microglia and astrocyte results might suggest that the effects are mainly caused by the combination treatment, however, differences between region-specific glucose uptake and brain inflammation implicates additional currently unknown mechanisms.

While trametinib and the combination treatment increased lifespan in both male and female mice, we observed significant sex-specific effect of the drug treatments especially on the molecular level. This includes downregulation of plasma cytokines, which was more prominent in female mice and the tissue-specific response in the transcriptome of aged animals. Sex-specific effects are commonly observed in longevity interventions [86] and have also been shown to occur in interventions targeting the insulin/mTOR network [87][88]. The reason for the sex-specific molecular response towards the drug treatments is currently unclear, but might reflect differences in the ageing process between male and female animals, or differences in the bioavailability of the drugs. Indeed, trametinib plasma levels were higher in female than male mice. Rapamycin blood levels were also higher in female UM-Het3 mice [28], but there was no sex difference in rapamycin plasma levels of the intermittently fed C3B6F1 mice used in this study [30]. Noteworthy, despite the higher plasma levels, the transcriptional response towards trametinib treatment was not necessarily stronger in female than male mice, which was especially obvious in the spleen, suggesting that other processes than just drug bioavailability contribute to the sexual dimorphic molecular response.

In summary, our results establish the FDA-approved drug trametinib as a new gero-protector in mammals that works additively with rapamycin to extend life and health span of mice. Trametinib at low doses is well tolerated in patients with only minor side effects and should therefore be included in future trials to address whether it also provides anti-ageing benefits in humans, especially in combination with rapamycin.

## Methods

### Mouse Husbandry and drug treatments

All animal protocols were done in accordance with the guidelines of the Federation of the European Laboratory Animal Science Association (FELASA) and approved by the Landesamt für Natur, Umwelt und Verbraucherschutz, Nordrhein-Westfalen, Germany (reference no. 84-02.04.2017.A074 and 81-02.04.2019.A313).

Mice were housed in individually ventilated cages (GM500 Mouse IVC Green Line, Techniplast), in groups of five per cage, under specific-pathogen-free conditions, at 21°C, with 50-60% humidity and a 12h light/dark cycle. Mice had constant access to nesting material and chew sticks and ad libitum access to a standard rodent diet (Ssniff, R/MH low phytoestrogen content, Ssniff Spezialdiäten GmbH) and sterile-filtered water. All mice were C3B6F1 hybrid, generated in-house by crossing C3H/HeOuJ females with C57BL/6NCrl males (strain codes 626 and 027, respectively, Charles River Laboratories). Pups were weaned at 3 weeks of age. Females were randomly allocated to cages. Males were weaned litter-wise to avoid aggression. If males of different litters had to be combined, a ratio of 2:3 was preferred over 4:1. Cages were randomly assigned to treatment groups, to which researchers were blinded. For the trametinib pre-test experiment, 3-month-old males and females, 5 per treatment group, were fed with standard chow containing 0.29, 0.58, 1.44, 2.88 or 11.52 mg trametinib / kg of diet and PEG-400 (3,2 ml/kg) for 4 weeks. Mice were sacrificed and tissues were dissected for molecular and pathological analyses. At weaning mice were fed a standard chow diet supplemented with the rapamycin encapsulation material Eudragit S100 (1-207-490-4242, Evonik CYRO LLC, 480 mg/kg diet). All experimental drug treatments started at 6 months of age, when control mice received a standard diet supplemented with the corresponding amount of Eudragit S100 (480 mg/kg) and PEG-400 (3,2 ml/kg). Animals treated with rapamycin received standard chow containing 42 mg/kg rapamycin (522 mg encapsulated rapamycin/kg) and PEG-400 (3,2 ml/kg) and animals treated with trametinib received food containing 1,44 mg/kg trametinib, Eudragit S100 (480 mg/kg) and PEG-400 (3,2 ml/kg). Animals treated with the combined rapamycin/trametinib diet received food containing 42 mg/kg rapamycin, 1.44 mg/kg trametinib and PEG-400 (3,2 ml/kg). Trametinib was administered continuously. Rapamycin, whether alone or with rapamycin, was administered alternating between rapamycin and control food on a weekly basis [30]. Rapamycin was obtained from LC Laboratories and encapsulated by Southwest Research Institute (SwRI). Trametinib was obtained from Biozol (1187431-43-1).

### Survival analysis

Survival of animals from six months on was assessed for a total of 878 mice, 484 males and 394 females (File 1). Mice for phenotyping, tissue collection and survival analysis were bred in three generations from the same breeding pairs. Mice of the phenotyping and tissue collection cohorts were included in the survival analysis and were censored at 24 months of age, when their issues were collected. Health status of all mice, including early signs of tumour growth, was monitored daily using a standardised health score, animals with poor health status were closely monitored and weighed daily and euthanized if they reached a predefined score. Mice were otherwise left undisturbed until they died of natural cause. Kaplan-Meier survival curves were generated using birth and death dates. Median lifespan was assessed, and survivorship was analysed using log rank test and Cox proportional hazard analysis. Maximum lifespan was assessed using Wang–Allison test. The survival curves for control and rapamycin treated mice were previously published [30].

### Tissue collection

Mice were sacrificed using cervical dislocation. Tissues were dissected and snap-frozen in liquid nitrogen and stored at -80°C. Plasma was isolated by adding 1μl of 500 mM EDTA per 100 μl of blood and subsequent centrifugation at 1,000g for 10 min at 4 °C. The supernatant was transferred to a fresh vial and stored at −80 °C. Tissues for histopathological analysis were fixed in 10% neutral-buffered formalin for preparation of formalin-fixed paraffin embedded (FFPE) tissues. Mice were sacrificed using cervical dislocation. Tissues were dissected and snap-frozen in liquid nitrogen and stored at -80°C. Plasma was isolated by adding 1μl of 500 mM EDTA per 100 μl of blood and subsequent centrifugation at 1,000g for 10 min at 4 °C. The supernatant was transferred to a fresh vial and stored at −80 °C. Tissues for histopathological analysis were fixed in 10% neutral-buffered formalin for preparation of formalin-fixed paraffin embedded (FFPE) tissues.

### Cross-sectional histopathology

Cross-sectional histopathology of liver, heart, pancreas, kidney, WAT, BAT, spleen, ovaries and testes of 24-month-old animals was carried out by Prof. Robert Klopfleisch (Institute of Veterinary Pathology, Freie Universität Berlin). Hematoxylin and eosin stainings were scored. Liver tissue was examined for the presence of lymphomas, sarcomas, benign or malignant tumours, lipidosis (diffuse or focal) and necrosis. Liver tumour grade was determined as 1 (low), 2 (moderate), 3 (high) and 4 (very high). Heart tissue was assessed for hypertrophy and fibrosis. In pancreas, inflammation, fat necrosis, lymphoma and exocrine atrophy were evaluated. For heart and pancreas, presence pathology was scored as 1 and absence as 0. In kidney, lymphoma, oxalate crystals and hydronephrosis were documented and kidney inflammation and glomerulopathy were scored with 1 (mild), 2 (moderate) and 3 (severe). WAT and BAT were scored for lymphoma, sarcoma, necrosis and granuloma and inflammation as 1, absence of pathology as 0. In spleen, extramedullary hematopoiesis, lymphomas, sarcomas, germinal centre and mantle zone hyperplasia were scored with 1 or 0 (absence). Testes were examined for atrophy or degeneration and ovaries for the presence of cysts, lymphoma and endometrial hyperplasia.

### Post-mortem macropathology

Both mice that died naturally and those sacrificed due to bad health were assessed for tumour load and location, organ enlargement and abnormalities, including organ discoloration or granulation. The presence and severity of tumours was scored with 0 (absence), 1 (one tumour in one organ), 2 (multiple tumours or tumour diameter > 3 cm in one organ or metastasis in two organs), 3 (severe metastasis in 3 or more organs) [89]. Other lesions were scored with 1 or 0 (absence). The severity of pathology was adjusted upon combination of multiple pathological findings (i.e. if one organ was affected by both tumours and discoloration, the score was adjusted from 1 to 2) [89].

### LC-MS analysis of trametinib plasma levels

20 uL of plasma was thawed on ice and mixed with 1 ml of a -20 °C methyl-tert-butyl-ether (MTBE):methanol:water (5:3:2 (v:v:v)) mixture, containing 10 ng/mL of everolimus as an internal standard. Samples were vortexed and incubated for 30 min at 4 °C. Protein pellets were obtained by centrifugation at 21.000 x g and 4°C for 10 min. Supernatant was transferred to fresh tubes and separation of the polar and lipid-containing phases facilitated by 150 μl of MTBE and 100 µL of UPC/MS-grade water (Biosolve). Samples were vortexed and incubated for 15 min at 15 °C. Phase separation was performed by 5 min centrifugation at 16.000×*g* at 15°C. 600 μl of the MTBE phase, containing the hydrophobic compounds (including everolimus and trametinib), was transferred to fresh microcentrifuge tubes. Extracts were dried in a Speed Vacuum concentrator at 30°C, and resuspended in 80µl of 70:30 [v:v] acetonitrile:isopropanol (Biosolve, Valkenswaard, Netherlands) containing 5% dimethylsulfoxid. Samples were vortexed and incubated for 5 min in a sonication bath and subsequently filtered through a 0.2µm filter. Resuspended samples were transferred to autosampler vials and liquid chromatography–mass spectrometry (LC-MS) performed using a Xevo-TQs triple quadrupole mass spectrometer (Waters) coupled to an Acquity iClass liquid chromatography system (Waters). The LC was equipped with a BEH C18 column (100 x 2.1 mm, Waters). Mobile phase A was UPLC-grade water (Biosolve), containing 10 mM ammonium acetate and 0.1% acetic acid (Sigma Aldrich). Mobile phase B contained acetonitrile: isopropanol 70:30 [v:v] (Biosolve). LC-based separation was performed using a 6 min gradient starting with 300 µL/min of 45% mobile phase B. This flow was held constant for 0.5 min, before ramping to 100% mobile phase B within the next minute. Composition was constant for the next 2 min, then ramped down within 0.1 min to the starting conditions of 45% mobile phase B. The column was equilibrated for 2.4 min (45% mobile phase B), before the next samples could be injected. The MS parameters were set to: Capillary voltage 1.5 kV, desolvation temperature 550°C, desolvation gas flow 800 l/h, cone gas flow 50 L/h and collision cell gas flow 9 L/h. Samples were analysed using a multiple reaction monitoring (MRM) method. For the absolute quantification of trametinib levels, trametinib standards were diluted to concentrations of 0, 2, 4, 10, 40, 200, 600, 1200, 2400 and 3600 ng/mL. The LC-MS data was analysed using the TargetLynx software (Version 4.1, Waters).

### Measurement of AST and ALP levels

Plasma measurements of AST, and ALP levels were performed by Laboklin GmbH & Co. KG, Diagnostic Laboratory, Bad Kissingen, Germany.

### Plasma proteomics (Olink)

The relative protein levels of 92 selected proteins were measured via proximity extension assay in EDTA plasma samples using the Olink Target 96 Mouse Exploratory panel (Olink proteomics, Uppsala, Sweden). Protein expression values were quantified by qPCR, measuring the amplification generated by oligonucleotide labelled antibody probe pairs bound to their respective target protein. After sample quality control by evaluation of internal detection controls, protein expression values were normalised using the Ct values of extension controls and a relative correction factor determined by Olink. Normalised protein expression values (NPX) on a log2 scale were analysed. 84 out of 86 plasma samples provided high quality data. After quality control, a total of 74 proteins were evaluated, representing >75% quantification values as determined by values above the limit of detection. Individual protein measurements at old age between treatment groups were compared using Kruskall-Wallis and post-hoc Mann Whitney U-test with Benjamini-Hochberg correction (scipy Python-library v1.6.2).

### Protein extraction for Western blot

5-10mg of liver tissue was lysed with 400 μl of RIPA buffer (Pierce) containing Complete mini protease inhibitor without EDTA (Roche) and PhosStop phosphatase inhibitors (Roche) in precooled FastPrep™ Lysing Matrix D tubes (MP Biomedicals™, 116913100) using FastPrep-24 (ThermoFisher Scientific, 4×25Hz, 30 sec). Protein extracts were incubated on ice for 10 min and placed in a sonicator water bath filled with ice for 10 min. Centrifugation for 15 min at 13.000 rpm and 4°C followed. Protein quantification was performed using the BCA Protein Assay Kit (Pierce™, 23225). Samples were boiled, Laemmli buffer at ¼ of the total volume and 5% β-mercaptoethanol were added and protein extracts were stored at -80°C.

### Western Blotting

20 μg of protein extracts were loaded and separated using 12% acrylamide gels (Criterion™ TGX Stain-Free™ Protein Gel (Biorad, 5678044) and blotted on PVDF membranes (AmershamTM Hybond, GE10600023, Merck) at 80V, for 1 hour, on ice. Membranes were blocked using 5% non-fat dry milk powder in Tris-buffered saline (TBS) 0.1% Tween 20 (TBS-T) at room temperature (RT) for at least 1 hour, then incubated with primary antibodies for phosphorylated ERK1/2 (1:1000, anti-rabbit, Cell Signaling Technology, 4370) and total ERK1/2 (1:1000, anti-rabbit, Cell Signaling Technology, 4695) in 5% fatty-acid free BSA in TBS-T at 4°C overnight. Blots were washed 3×15 min in TBS-T and incubated with HRP-coupled secondary antibodies (ThermoFisher) diluted in 5% milk in TBS-T for 1h at RT, followed by 3×15 min washes in TBS-T at RT. For signal development, ECL Select Western Blotting Detection Reagent (GE Healthcare) was applied and image acquisition was performed using a ChemiDoc™ XRS+ System (Bio-Rad). Western Blot signals were quantified using the Image Lab™ Software (Bio-Rad) with alpha tubulin (1:1000, anti-rabbit, Cell Signaling Technology, 2125) as internal control and normalised against non-drug treated controls.

### Immunostaining of brains

Brains of 6-month-old control females and 24-month-old control, rapamycin, trametinib and rapamycin/trametinib females were dissected and cut in half to facilitate fixative penetration in 10% neutral-buffered formalin for 2-4 hours at RT. Brains were fixed overnight at 4°C, then dehydrated in 30% sucrose in 1x phosphate-buffered saline (PBS) for 2-4 hours at RT and overnight. Following dehydration, brains were frozen in Tissue-Tek OCT (Labtech) on dry ice and stored at −80 °C. Brains were cut using a cryostat (CM1850, Leica Biosystems). 25-μm, sections were mounted on Superfrost Plus microscope slides (ThermoFisher Scientific) and stored at −80 °C. For immunostainings, slides were washed with PBS for 10 min, permeabilized with PBS + 0.2% Triton X-100 (PBST) for 15 min and washed in PBST for 5 min. Samples were incubated with 2% normal goat serum in PBST at RT and then incubated with primary antibodies in 2% normal goat serum in PBST overnight at 4 °C. Primary antibodies were: Ionised calcium-binding adaptor molecule 1 (Iba-1, 1:500, 019-19741, Wako) and Glial fibrillary acidic protein (GFAP, 1:1000, G3893, Sigma). Slides were washed 3×15 min in PBST and incubated for one hour with Alexa Fluor-conjugated secondary antibodies at 1:1000 in 2% normal goat serum at RT. Secondary antibodies were: Alexa Fluor 488-goat anti-rabbit IgG (A-11008, to detect Iba-1) and Alexa Fluor 633-goat anti-mouse IgG (A-21126, to detect GFAP). Slides were washed 2×10 min in PBST and 1×10 min in PBS and incubated for 30 min with DAPI (1:10000 in PBS). Slides were washed with PBS for 10 min and embedded using VectaShield Antifade Mounting Medium without DAPI (Vectorlabs).

### Confocal imaging and quantification of astrocyte and microglia density

Imaging was performed using a Leica SP8-Upright confocal microscope (Leica Microsystems). HyD detectors for Alexa Fluor 488 and 663 with 6% and 10% power respectively and the corresponding gating and the excitation wavelength settings to maximise fluorophore emission were used. 40-μm *z*-stack confocal images were acquired at 2-μm intervals, with 40×/1.3 oil objective at 1× zoom. Settings were standardised for all treatment groups during image acquisition. Representative montages from *z*-stack confocal images were used and image processing was performed using Image J (Fiji) software v2.3.0/1.53q (Image J, Maryland, USA). For quantification of astrocytes and microglia density, the cell counter tool was used to count GFAP+ or Iba-1+ cells that overlapped with DAPI+ nuclei, respectively in 10-15 images per brain region per mouse brain.

### RNA isolation

4-10 mg of spleen, muscle and kidney samples were homogenised in 1 mL TRIzol (Life Technologies) using a FastPrep-24 (MP Biomedicals) with 6 times bead-beating at 4 m/s for 30 sec. Samples were incubated at RT for 5 min, 200 μl of chloroform was added and mixed by vortexing, and samples incubated at RT for 10 min and centrifuged at 12000 g for 15 min at 4 °C. The aqueous upper phase was collected and transferred to a new RNAse-free tube. 500 μl of isopropanol, 50 μl of 3.0 M NaOAc and 1.5 μl of glycogen (ThermoFisher Scientific) were added,mixed and incubated at -80 °C for 30 min. Samples were centrifuged at 12.000 g for 10 min at 4 °C. RNA pellets were washed twice with 1 ml of ice-cold 70% ethanol, air-dried for 5-10 min and then resuspended in 30 μl RNAse free water. To remove genomic DNA, RNA samples were treated with DNAse using the TURBO DNAse kit (ThermoFisher Scientific) according to the manufacturer instructions.

### RNA seq analysis

RNA-seq library preparation and sequencing (50 million reads, 25 million in each direction, 150bp paired end) of kidney, spleen and muscle RNA was performed by Admera Health (New Jersey, USA). poly(A)+ mRNA was isolated using the NEBNext Poly(A) mRNA Magnetic Isolation kit (New England BioLabs Inc., Massachusetts, USA). Prior to first strand synthesis, samples were randomly primed (5’ d(N6) 3’ [N=A,C,G,T]) and fragmented. First strand synthesis was done using the Protoscript II Reverse Transcriptase for approximately 40 minutes at 42⁰C. All remaining steps for library construction were done according to the NEBNext Ultra™ II Non-Directional RNA Library Prep Kit for Illumina (New England BioLabs Inc., Massachusetts, USA). Final libraries quantity was assessed by Qubit 2.0 (ThermoFisher, Massachusetts, USA) and quality was assessed by TapeStation D1000 ScreenTape (Agilent Technologies Inc., California, USA). The final library size was about 430bp with an insert size of about 300bp. Illumina 8-nt dual-indices were used. Equimolar pooling of libraries was performed based on QC values and sequenced on an Illumina Novaseq S4 (Illumina, California, USA) with a read length configuration of 150 bp for 50 M paired-end reads per sample, i.e. 25M reads for each direction). Libraries read processing was automated using Flaski (v3.11.34) [90], developed by the Bioinformatics Core Facility at the MPI for Biology of Ageing (Cologne, Germany). Raw reads were mapped to the mm39 ENSEMBL build 105 using kallisto (v0.46.1) [91]. Gene counts were quantified in the same step. Differential gene expression was analysed using DESeq2 (v1.24.0) [92] in R (v3.6.3). Genes were annotated as expressed if they were contained in the DESeq2 output. Differentially regulated genes (adjusted *p* value < 0.05) were additionally subsetted into total genes, upregulated genes (log2 fold change > 0), and downregulated genes (log2 fold change < 0). Data were processed using custom python scripts and the following packages: numpy (v 1.12.0), scipy (v 1.7.1), and pandas (v 1.2.0). Results were visualized using the following packages: matplotlib (v 3.3.2), matplotlib-venn (v 0.11.5), and seaborn (v 0.11.1). Gene ontology enrichment analysis was performed using the DAVID API based function DAVIDenrich of the AGEpy python package (v 0.8.2) on the GOTERM_BP_FAT ontology to identify significantly enriched terms (Benjamini-Hochberg adjusted *p*-value < 0.05). As background, all expressed genes of a tissue were used. Redundancy of significantly enriched GO terms (adjusted *p*-value < 0.05) was reduced using REVIGO (cutoff: “0.7”, valueType: “pvalue”, measure: “SIMREL”). GO terms were visualized using dotplot representation.

### 18F-FDG PET/CT Imaging

PET imaging was as described in [93] using an Inveon preclinical PET/CT system (Siemens). Mice were anesthetized with 2% isoflurane in 65%/35% nitrous oxide/oxygen gas and positioned on a dedicated mouse carrier (MEDRES, Germany) carrying two mice. For injection of the radiotracer, a catheter consisting of a 30G cannula connected to a polythene tubing (ID = 0.28 mm) was inserted into the tail vein and fixated by a drop of glue. After starting the PET scan, 7-8 MBq of [18F]-FDG in 50-100 mL saline were injected per mouse. Emission data were acquired for 45 minutes. Thereafter, animals were automatically moved into the CT gantry and a CT scan was performed (180 projections/360°, 200 ms, 80 kV, 500 mA). The CT data were used for attenuation correction of the PET data and the CT image of the scull was used for image co-registration. Plasma glucose levels were determined from a tail vein blood sample using a standard glucometer (Bayer) after removing the tail vein catheters. PET data were histogrammed in time frames of 12×30s, 3×60s, 3×120s, 7×240s, rebinned in 3D, and images were reconstructed using the MAP-SP algorithm provided by the manufacturer. The imaging analysis software Vinci [94] was used to co-register images to a 3D mouse brain atlas constructed from the 2D mouse brain atlas published by Paxinos [95]. An image-derived input function was extracted from the PET data of the aorta, which could be identified in the image of the first-time frame of each animal. Input function data were corrected for partial volume effect by assuming a standardized volume fraction of 0.6 [96]. Parametric images of the [18F]FDG kinetic constants K1, k2, k3, and k4 were determined by a voxel-by-voxel (voxel size = 0.4 mm x 0.4 mm x 0.8 mm) fitting of data to a two-tissue-compartment kinetic model. The ratio of tissue and plasma glucose concentrations (CE/CP) is a measure for glucose transport and is given by CE/CP = K1/(k2+k3/0.26) [35] [93]. Since cellular activation is accompanied by increased glucose transport and this parameter is less sensitive to changes in plasma glucose level, we use alterations of glucose transport (CE/CP) as surrogate for alterations in neuronal activation. Statistical testing was performed by application of a voxel-wise t test between groups. 3D maps of p values allow for identification of regions with significant differences in the parameters. From these regions we defined volumes of interest (VOIs) and performed additional statistical testing.

### Mouse phenotyping

A separate mouse cohort of 15 mice peer treatment group was used to evaluate drug and age-related changes in health span, behaviour and fitness. Experiments were performed at 12 (middle-aged) and 20-22 (old) months of age. All tests were carried out during the light phase and experimenters were blinded to the treatment group.

### Electrocardiography

Electrocardiography (ECG) was performed non-invasively using the ECGenie system (Mouse Specifics). For acclimatisation, each mouse was placed on the training platform for 10 minutes and then placed on the measurement platform on interchangeable electrodes that transmit the electrical signals to a computer for a maximum of 10 minutes. Signals with a heart rate variation that exceeded 35 bpm were excluded [97] and average heart rate per mouse was evaluated.

### Rotarod

Rotarod was performed to assess motor coordination using the Rotarod (TSE Systems, type 3375-M5). Measurement started at low speed (5 rpm) and the revolving rate was continuously increased to 40 rpm over a period of 300 seconds. The total time that each mouse spent on the rod, with a cut-off time of 300 seconds was measured. The test was performed on four consecutive days and only data from day four were used for the analysis.

### Treadmill

To measure endurance, mice were placed onto the treadmill (TSE Systems, type 3033401-M-04/C) and acclimatised to the experimental setup for 5 minutes under low speed (0.1m/sec). A progressively accelerating speed was applied from 0.1 m/sec for another ten minutes to 1.3 m/s within 60 min. A mild electroshock (0.3 mA) was applied for 5 seconds, followed by 5 seconds of refractory period as soon as mice slowed down beyond a critical point and crossed the laser beam for more than 2 seconds to ensure that mice only stopped running upon exhaustion. Exhaustion was defined as the willingness of a mouse to withstand three consecutive shocks instead of continuing running.

### Open field

The open field experiment was carried out to determine locomotion, exploratory drive and anxiety-like behaviour (ActiMot2, TSE Systems). Mice were placed in a 50 cm x 50 cm x 40 cm box and were allowed to freely explore for 10 minutes. The test chamber was illuminated to 20-30 lux. Total activity, speed and time spent in the centre was recorded via infrared sensors.

### Y maze

Mice were placed in a Y-shaped maze and their activity recorded for 5 minutes using a video tracking system (VideoMot, TSE Systems). Global speed indicates the average speed of the mice during the 5 min measurement period. % alternating arm visits were calculated as a read out for spatial working memory.

### Statistical analysis

Experiments were performed in a randomised and blinded fashion whenever possible. Statistical analysis was performed using GraphPad Prism 9.0, except for Cox Proportional Hazard analysis which was performed in R (R Core Team). Number of animals and statistical tests are indicated in the Figs/Fig legends. For multiple comparison testing, One-way and two-way Analysis of variance (ANOVA) were followed by Bonferroni post-hoc test. To test for differences in the proportion of mice affected by pathologies, Chi Square test and Poisson regressions were used. The counts of mice where the pathological finding was present or absent was set as the dependent variable and the absence of pathology (score 0) was set as the reference classification. Comparisons were performed against control mice as reference. Error bars indicate standard error of mean (SEM). Significance is indicated as: *P < 0.05; **P < 0.01; ***P < 0.001; ****P < 0.0001.

## Data availability

All data that support the findings of this study are available from the corresponding authors upon reasonable request.

## Acknowledgements

We thank Sarah Veugelen and Maarouf Baghdadi for their help in establishing the mouse brain stainings. We acknowledge Andrea Mesaros and Martin Purrio from the Phenotyping facilitiy of the Max Planck Institute for the Biology of Ageing for outstanding technical help and advice. We are very grateful to the Comparative Biology facility of the Max Planck Institute for Biology of Ageing for mouse housing. Oliver Hendrich, Sandra Buschbaum, André Pahl, Jenny Fröhlich and Ramona Jansen are acknowledged for technical assistance with mouse experiments. Imaging was performed in the FACS & Imaging Core Facility at the Max Planck Institute for Biology of Ageing.

## Contributions

Conceptualization of the study was carried out by T.N., S.G. and L.P. Experiments were conducted by L.G., A.L.C., H.B. and T.N. J.B. and H.B. conceived the PET experiment. A.L.C. and H.B. performed and analysed the PET experiment. C.M. analysed the O link measurements. T.N. analysed the RNA seq data. A.B. supervised the RNA seq analysis. L.G. performed brain stainings. Statistical evaluation of data was done by LG, C.M., H.B., T.N. and S.G. Writing of the manuscript by L.G., S.G. and L.P.

## Corresponding author

Correspondence to Linda Partridge or Sebastian Grönke

## Ethics declarations Competing interests

The authors declare no competing interests.

## Supplemental Figures

**Sup Figure 1.**
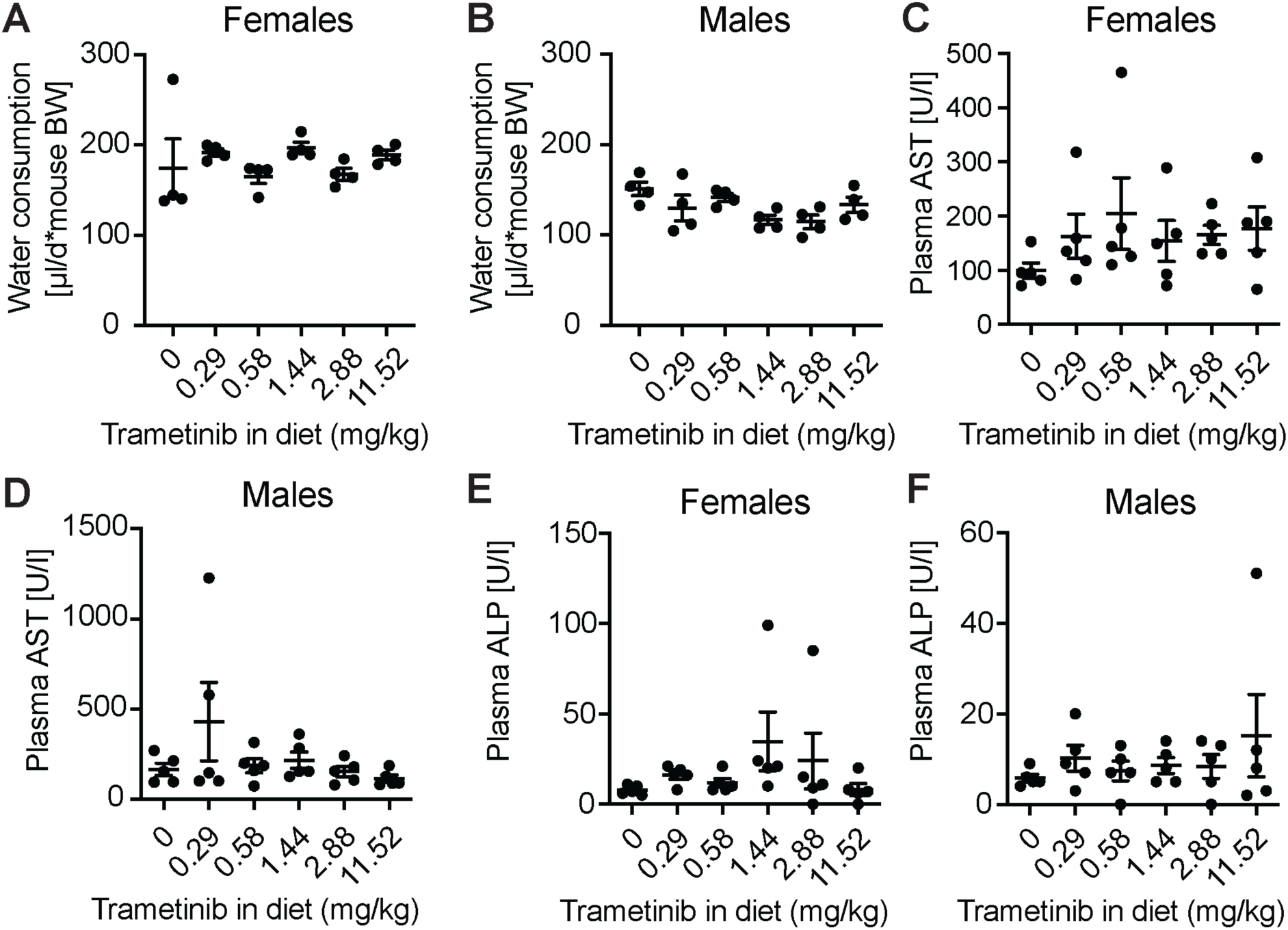
Water consumption and biomarkers of liver health are not affected by 4 weeks of trametinib treatment in each sex. **a**-**b** Water consumption of **a** females and **b** males were not significantly changed. Plasma level of **c**-**d** Aspartate transaminase (AST) and **e**-**f** Alkaline Phosphatase (ALP) as biomarkers of liver health were not significantly changed. Significance was tested by One-Way ANOVA. N=5 per treatment for both sexes.

**Sup Figure 2.**
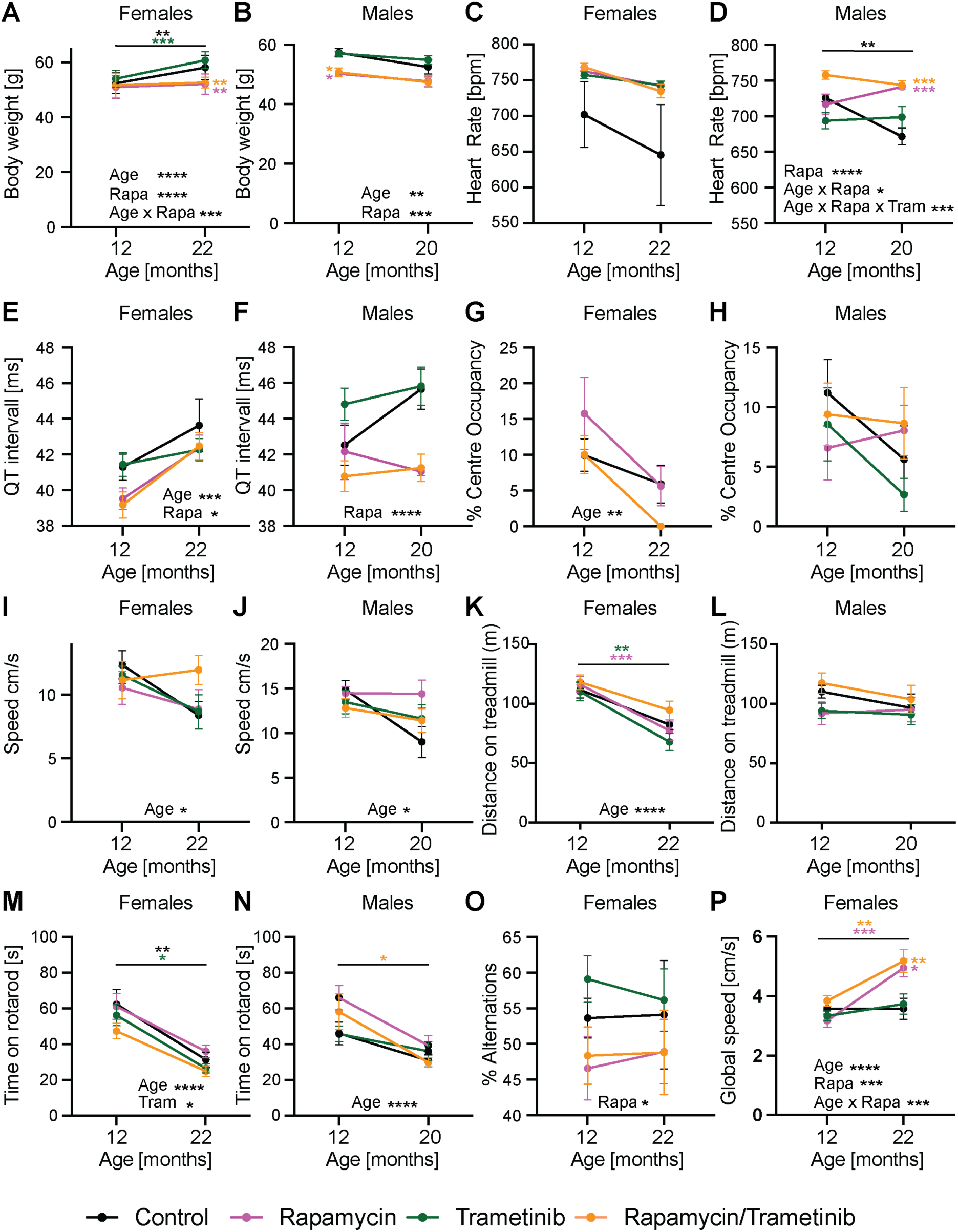
Rapamycin treatment reduces body weight, improves heart function and increases movement speed in old mice. **a-b** body weight, **c-f** ECG measurement of **c**-**d** heart rate and **e**-**f** QT-interval, **g**-**j** open field assay measurement of **g-h** centre occupancy and **i**-**j** movement speed, **k**-**l** distance on treadmill, **m**-**n** time on rotarod, **o-p** Y maze measurement of **o** % activity in new arm and **p** global speed. Data were analysed using a mixed effect model testing for the interaction between Rapamycin x Trametinib x Age followed by a Bonferroni post hoc test to test for significance changes between 12 and 22 months indicated by asterisks above the connecting lines and between drug treatments and control at 12 or 22 months indicated by asterisks next to the dots. *p<0.05, **p<0.01, ***p<0.001, ****p<0.0001. Significant terms from the interaction analysis and the corresponding p-value are indicated per panel. N numbers (12M, 22M) for: female body weight, rota rod: Control (15/13), Rapamycin (15/13), Trametinib (15/10) Rapamycin/Trametinib (15/15); male body weight, rota rod: Control (14/10), Rapamycin (15/13), Trametinib (15/13), Rapamycin/Trametinib (15/15); female heart rate, QT-interval: Control (15/9), Rapamycin (15/10), Trametinib (8/10) Rapamycin/Trametinib (7/7); male heart rate, QT-interval: Control (15/9), Rapamycin (15/10), Trametinib (8/10) Rapamycin/Trametinib (7/7). N numbers (22M female, 20M male) for: open field: Control (13,11), Rapamycin (14,13), Trametinib (12,13) Rapamycin/Trametinib (15,15); treadmill: Control (13,11), Rapamycin (14,13), Trametinib (12,13), (Rapamycin/Trametinib (15,15).

**Sup Figure 3.**
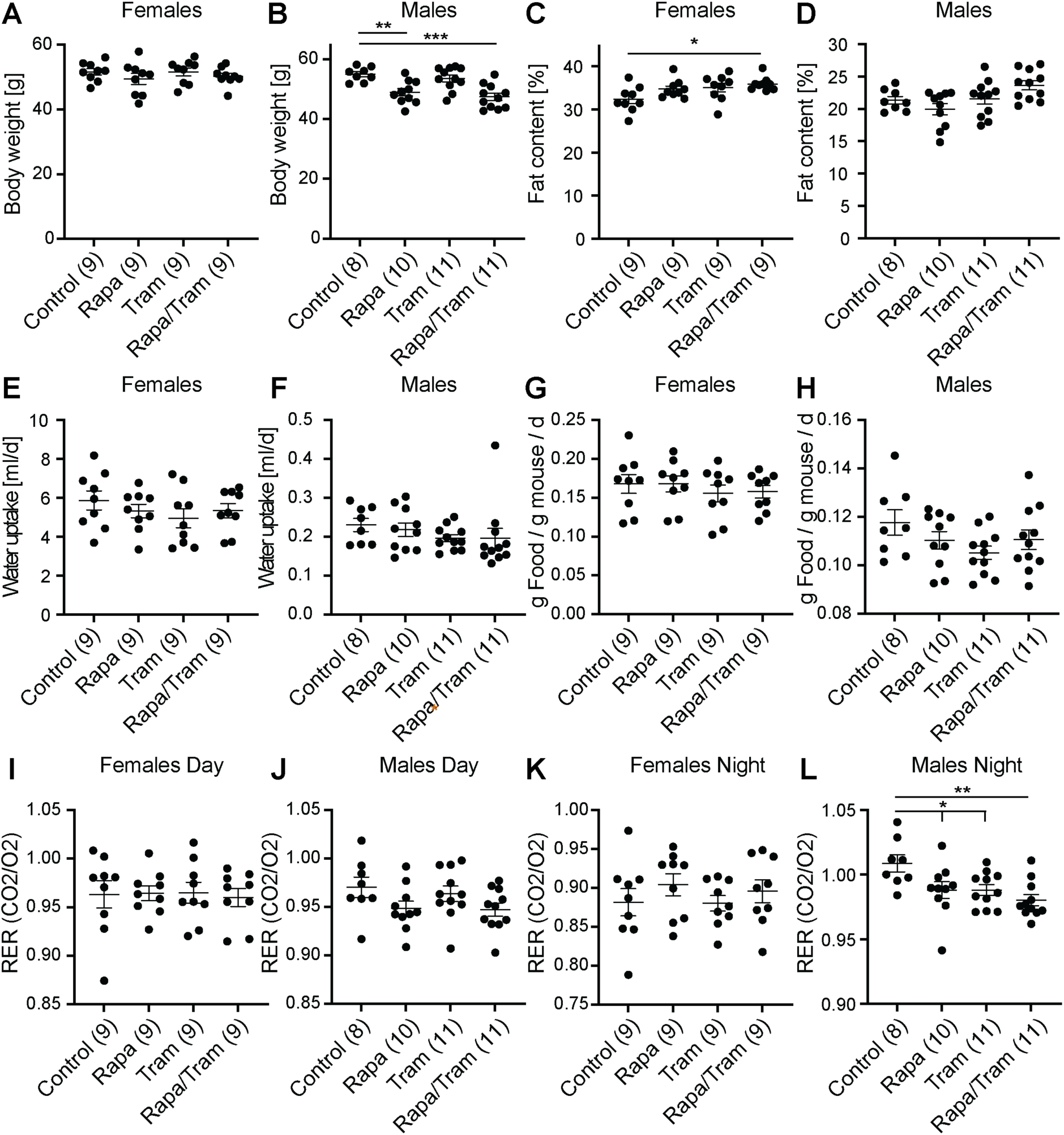
Rapamycin and trametinib do not affect food or water uptake but do affect night time metabolism in males. **a-b** body weight. **c**-**d** nuclear magnetic resonance measurement of % body fat. Metabolic cages were used to measure **e**-**f** water uptake, **g**-**h** food uptake relative to body weight and **i**-**l** respiratory exchange ratio (RER) during **i**-**j** day and **k**-**l** night. Measurements were done in batches between the age of 15-18 months. Rapamycin and combined treatment significantly reduced body weight in male but not female mice, but had no effect on water or food uptake. Fat content was significantly increased upon combined treatment in females and in tendency in males when compared to control animals. All drug treatments significantly reduced night-time RER in males but not females. *p<0.05, **p<0.01, ***p<0.001. One-Way ANOVA with Bonferroni post hoc test. Numbers in brackets indicate the number of animals assessed per treatment. Data are presented as mean ± SEM.

**Sup Figure 4.**
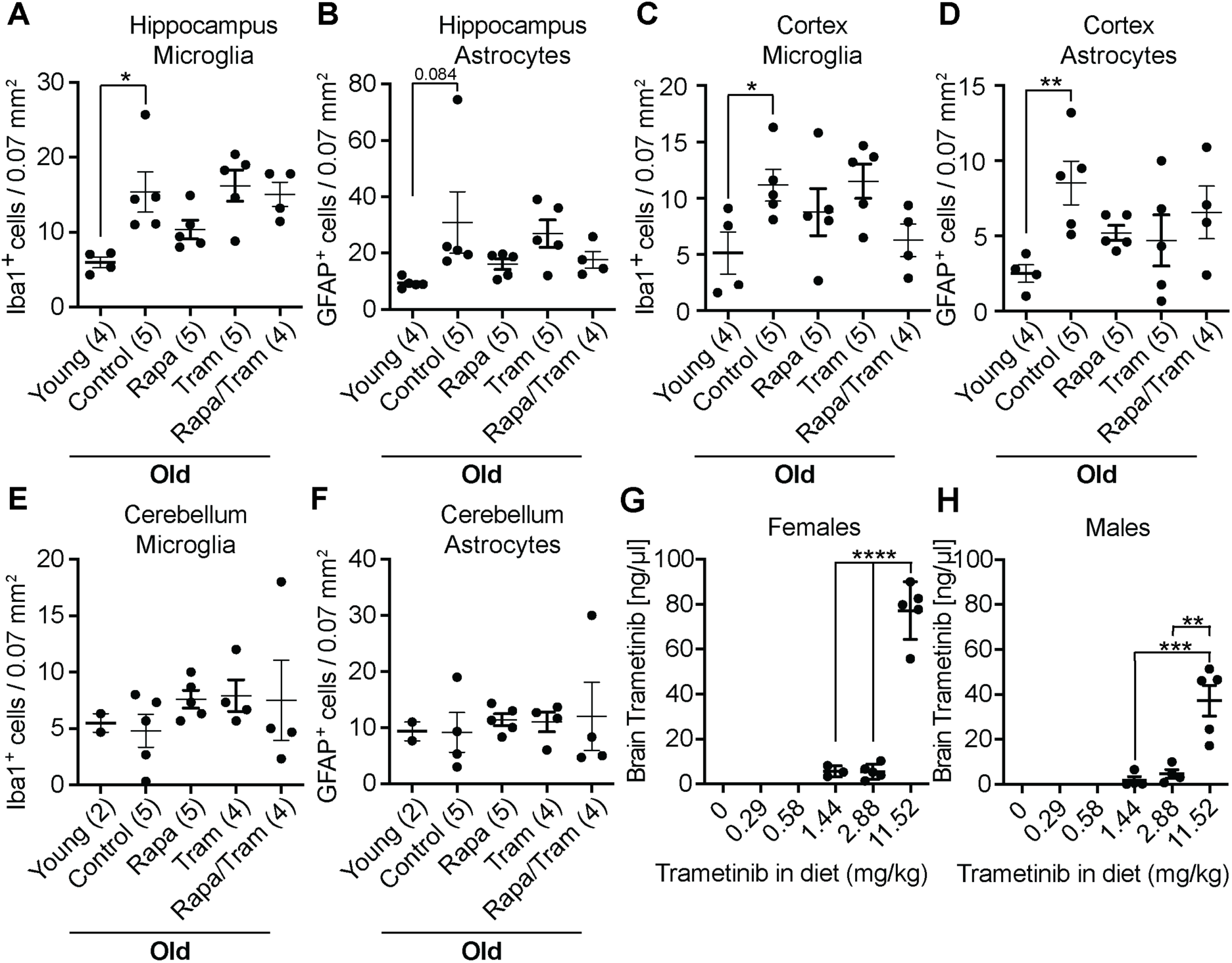
Combined rapamcin and rametinib treatment affects brain inflammation in a region-specific manner. Quantification of immunohistochemical stainings measuring the density of activated microglia (DAPI/Iba-1 positive cells) and astrocyte (DAPI/GFAP positive cells) in **a**-**b** hippocampus, **c**-**d** cortex and **e**-**f** cerebellum of 6-months old control animals (young) and 24 months-old control and drug-treated females. Microglia and astrocyte density significantly increased with age in the hippocampus and cortex of control animals, but not in the cerebellum. Single drug treatment did not significantly reduce microglia or astrocyte density in the hippocampus, cortex or cerebellum. **g**-**h** Brain levels of trametinib increased in a dose-dependent manner upon feeding **g** female or **h** male animals for 4 weeks with food containing 0, 0.29, 0.58, 1.44, 2.88 or 11.52 mg/kg of drug. 5 brain samples were measured via mass spectrometry per treatment. Only samples with significant trametinib levels above background are shown. Trametinib was undetectable in the brain of animals fed with 0, 0.29 and 0.58 mg/kg trametinib, and only very low levels were detected in animals fed with 1.44 and 2.88 mg/kg trametinib. Only at 11.52 mg/kg trametinib, higher levels of trametinib were detected in the brain. *p<0.05, **p<0.01. One-Way ANOVA with Bonferroni post hoc test. 10-15 confocal images per brain region were averaged per sample in **a**-**g**. Numbers in brackets indicate the n number per treatment. Data are presented as mean ± SEM.

**Sup Figure 5.**
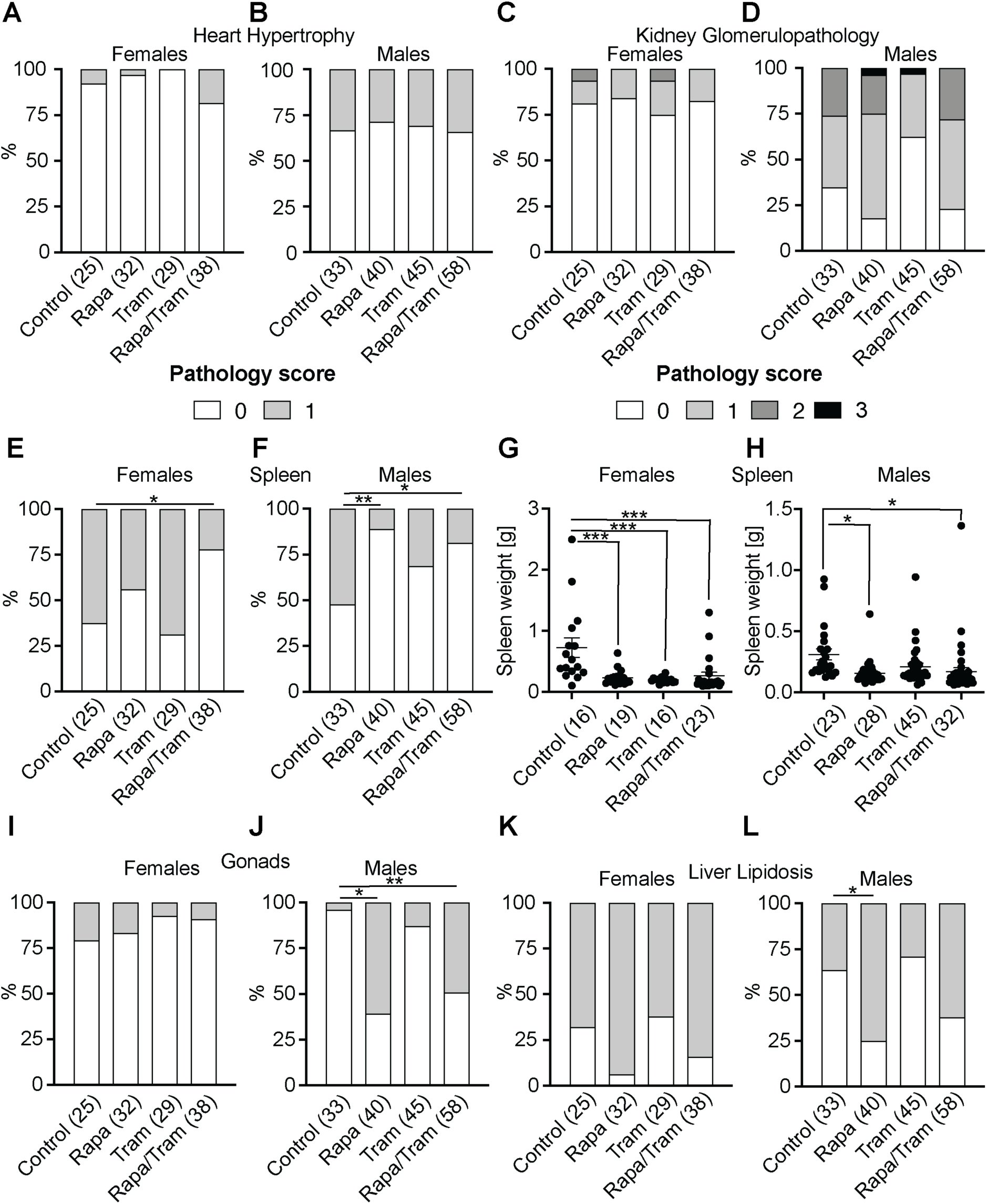
Combination of rapamycin and trametinib ameliorates spleen pathology but does not protect against rapamycin-induced liver lipidosis and testicular lesions. Cross sectional histopathological analysis of non-neoplastic lesions in tissues isolated from 24 months old male and female mice treated with rapamycin, trametinib and rapamycin/trametinib. **a**-**b** heart hypertrophy, **c**-**d** spleen pathology, **e**-**f** spleen weight, **g**-**h** kidney glomerulopathology, **i-j** gonadal pathology and **k**-**l** liver lipidosis. **a**-**b** Rapamycin and Trametinib treatments did not significantly affect heart hypertrophy. **c-d** Females and males treated with Rapamycin/Trametinib showed reduced spleen pathology and a similar trend was observed upon Rapamycin treatment, which was significant in males. **e**-**f** Spleen weight was significantly reduced upon Rapamycin and Rapamycin/Tram treatment in males and females and upon trametinib treatment in females. **g-h** Kidney glomerulopathology was not significantly affected by the drug treatments. **i-j** Males but not females treated with Rapamycin and Rapamycin/Trametinib showed a significant increase in gonadal pathology. **k**-**l** Male mice treated with Rapamycin showed a significant increase in liver lipidosis compared to control animals. A similar non-significant trend was also observed in female animals and upon Rapamycin/Trametinib treatment in males and females. Data in **a-d** and **g-l** are presented as percentage over total and statistical analysis was performed by Chi-square test and Poisson regression. Data in **e-f** are plotted as mean with SEM and statistical analysis was done using One-Way ANOVA followed by Bonferroni post hoc test. *p<0.05, **p<0.01, ***p<0.001. Number of scored tissues per treatment are indicated in brackets. Control and Rapamycin data in **a-l** were published previously (Baghdadi et al., 2024).

**Sup Figure 6.**
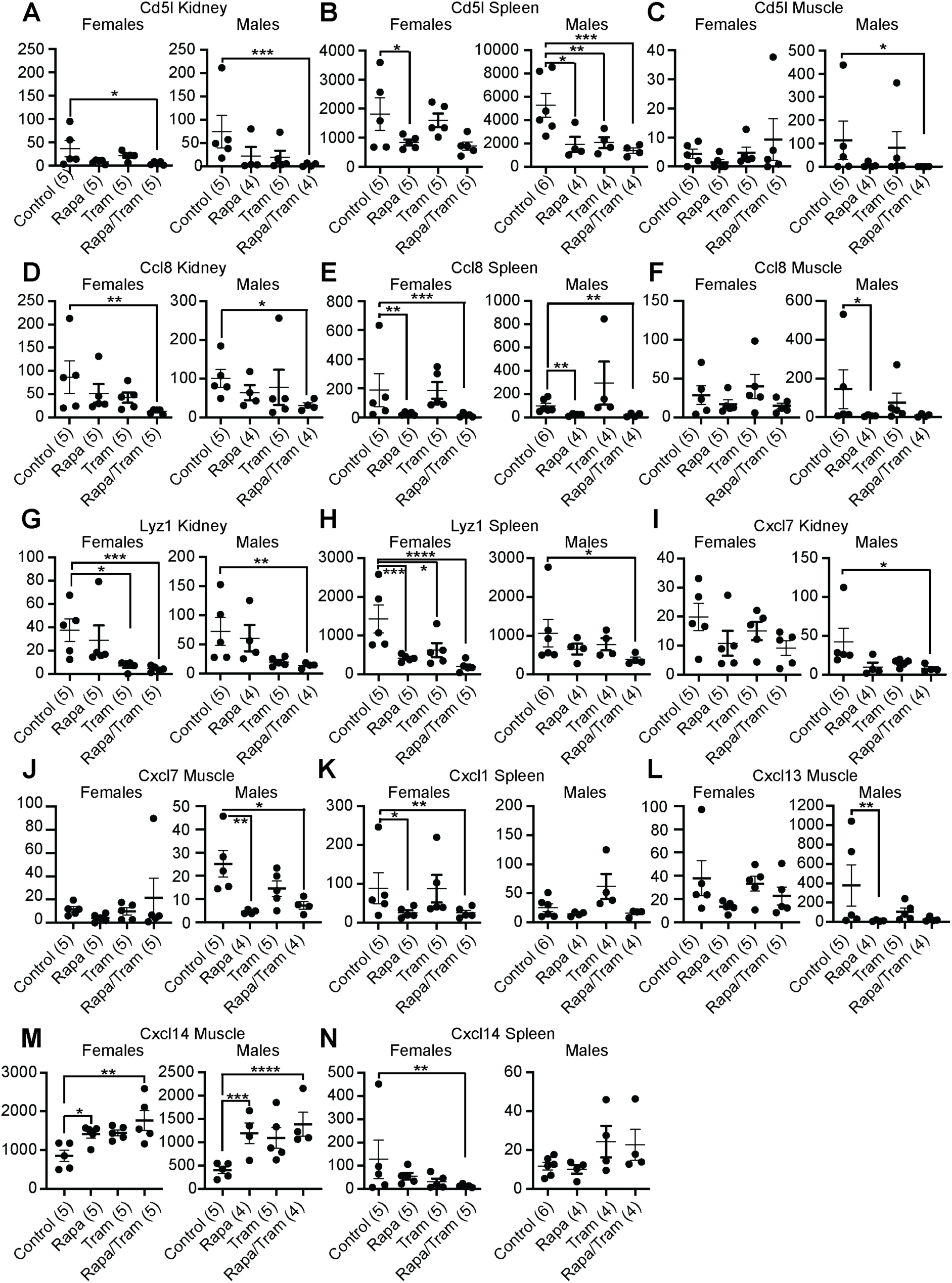
Genes involved in inflammation are downregulated upon combined rapamycin and trametinib treatment. Expression level of inflammation associated genes based on the RNA seq analysis of kidney, spleen and muscle tissue from 24-months-old rapamycin, trametinib and combination treated animals. **a-c** Cd5l was significantly downregulated in **a** kidney, **b** spleen and **c** muscle upon combination treatment. **d-e** Ccl8 was significantly downregulated by combination treatment in **d** kidney and by rapamcin and combination treatment in **e** spleen and **f** male muscle. **g-h** Lyzosyme1 expression was decreased in **g** kidney upon trametinib and combination treatment, and **h** in female spleen upon rapamycin, trametinib and combination treatment. **i-j** Cxcl7 was significantly downregulated in **i** male kidney and **j** male muscle upon combination treatment, and by rapamycin in male muscle. **k** Cxcl1 expression level was decreased in the female spleen upon rapamycin and combination treatment. **l** Cxcl13 was significantly downregulated by rapamycin in male muscle tissue. **m-n** Expression of Cxcl14 was upregulated by rapamycin and the combination treatment in **m** male and female muscle. **n** Cxcl14 was significantly downregulated in the female spleen. Data are presented as Mean ± SEM. Statistical analyses were performed using DESeq2/1.24.0. P-values were adjusted for multiple testing. *P < 0.05 and **P < 0.01, ***P < 0.001, and ****P < 0.0001. Number of scored tissues per treatment are indicated in brackets.

**Sup Figure 7.**
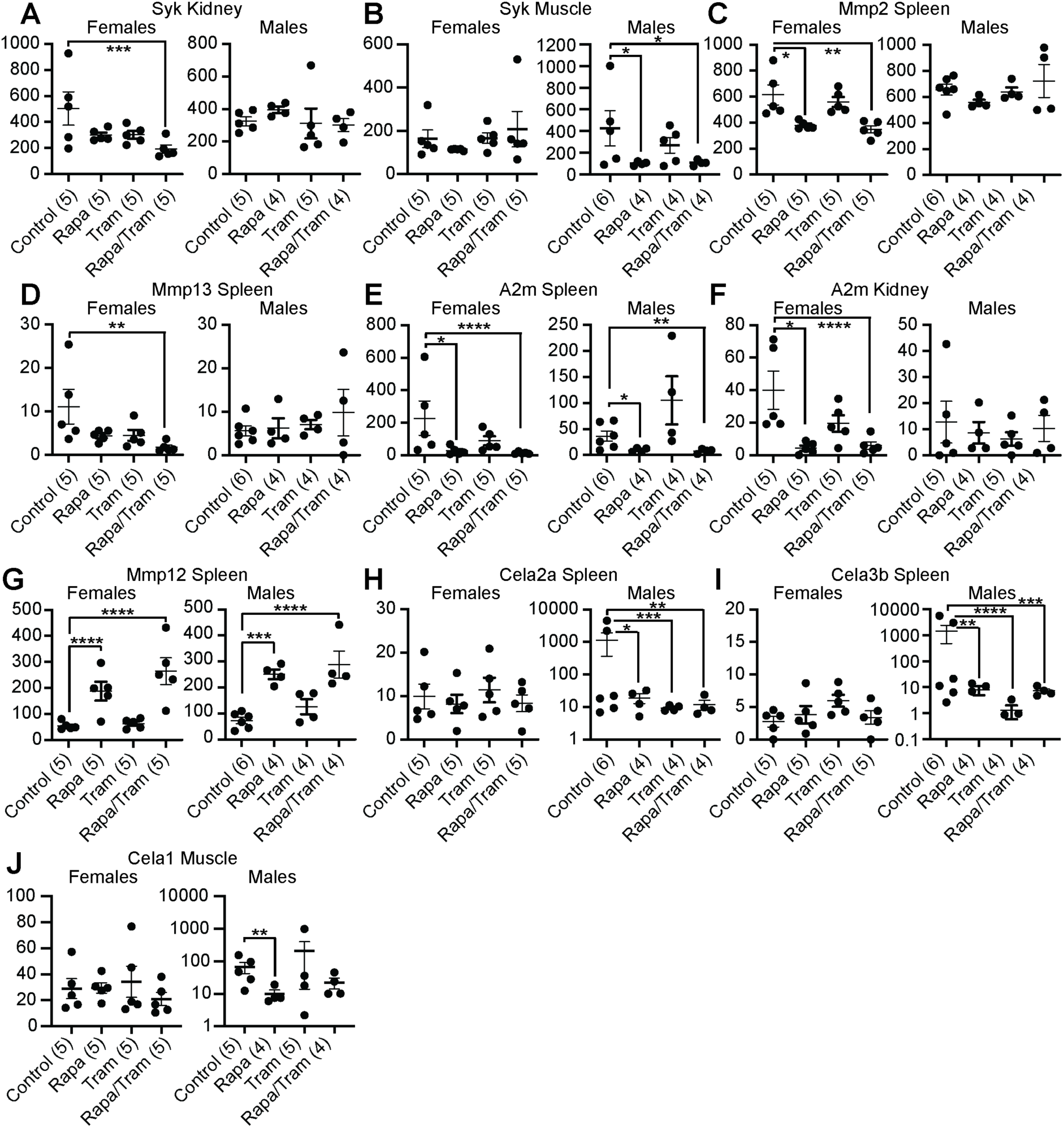
Ets2 target genes are downregulated upon combined rapamycin and trametinib treatment. Expression level of potential ETS target genes based on the RNA seq analysis of kidney, spleen and muscle tissue from of 24-months-old rapamycin, trametinib and combination treated animals. **a** Expression of SYK was significantly decreased in the kidney of female mice treated with rapamycin and trametinib, and **b** in male muscle upon rapamycin and combination treatment. **c** Mmp2 was downregulated in the female spleen upon rapamycin and combination treatment, **d** Mmp13 expression was decreased in the female spleen upon combination treatment. **e-f** A2m expression levels were decreased upon rapamycin and combination treatment in **e** male and female spleen and **f** female kidney. **g** Mmp12 levels were increased upon rapamycin and combination treatment in the spleen. **h** Cela2a and **i** Cela3b levels were specifically downregulated in male spleens upon single and combination drug treatment. **j** Cela1 expression was downregulated in male muscle upon rapamycin treatment. Data are presented as Mean ± SEM. Statistical analyses were performed using DESeq2/1.24.0. P-values were adjusted for multiple testing. *P < 0.05 and **P < 0.01, ***P < 0.001, and ****P < 0.0001. Number of scored tissues per treatment are indicated in brackets.

**Sup Table 1: Raw data of plasma proteins measured via the Olink Target 96 Mouse Exploratory panel in 24-months-old male and female mice.**

**Sup Table 2: Significantly regulated genes based on the RNA seq analysis in 24 months old mice treated with rapamycin, trametinib and the combination.**

